# Poly ADP-Ribose Signaling is Dysregulated in Huntington Disease

**DOI:** 10.1101/2022.11.23.517669

**Authors:** Tamara Maiuri, Carlos Barba Bazan, Rachel J. Harding, Nola Begeja, Tae-In Kam, Lauren M. Byrne, Filipe B. Rodrigues, Monica M. Warner, Kaitlyn Neuman, Muqtasid Mansoor, Mohsen Badiee, Morgan Dasovich, Keona Wang, Leslie M Thompson, Anthony K. L. Leung, Sara N. Andres, Edward J. Wild, Ted M. Dawson, Valina L. Dawson, Cheryl H. Arrowsmith, Ray Truant

**Affiliations:** Department of Biochemistry and Biomedical Sciences, McMaster University, Hamilton, ON L8S 3Z5, Canada; Structural Genomics Consortium, University of Toronto, Toronto, ON M5G 1L7, Canada; Department of Pharmacology and Toxicology, University of Toronto, Toronto, ON M5S 1A8, Canada; Neurodegeneration and Stem Cell Programs, Institute for Cell Engineering, Johns Hopkins University School of Medicine, Baltimore, MD 21205, USA; Department of Neurology, Johns Hopkins University School of Medicine, Baltimore, MD 21205, USA; UCL Huntington Disease Centre, UCL Queen Square Institute of Neurology, University College London, London, UK; Department of Biochemistry and Biomedical Sciences, McMaster University, Hamilton, ON L8S 3Z5, Canada; Michael G. DeGroote Institute for Infectious Disease Research, McMaster University, Hamilton, ON, Canada; Department of Biochemistry and Molecular Biology, Bloomberg School of Public Health, Johns Hopkins University, Baltimore, MD 21205, USA; Department of Neurobiology and Behavior, University of California, Irvine, CA 92697, USA; Department of Neurobiology and Behavior, University of California, Irvine, CA 92697, USA; Department of Psychiatry and Human Behavior, University of California, Irvine, CA 92868, USA; Department of Biochemistry and Molecular Biology, Bloomberg School of Public Health, Johns Hopkins University, Baltimore, MD 21205, USA; Department of Molecular Biology and Genetics, Department of Genetic Medicine, Department of Oncology, School of Medicine, Johns Hopkins University, Baltimore, MD 21205, USA; Neurodegeneration and Stem Cell Programs, Institute for Cell Engineering, Johns Hopkins University School of Medicine, Baltimore, MD 21205, USA; Department of Neurology, Johns Hopkins University School of Medicine, Baltimore, MD 21205, USA; Department of Pharmacology and Molecular Sciences, Johns Hopkins University School of Medicine, Baltimore, MD 21205, USA; Solomon H. Snyder Department of Neuroscience, Johns Hopkins University School of Medicine, Baltimore, MD 21205, USA; Neurodegeneration and Stem Cell Programs, Institute for Cell Engineering, Johns Hopkins University School of Medicine, Baltimore, MD 21205, USA; Department of Neurology, Johns Hopkins University School of Medicine, Baltimore, MD 21205, USA; Department of Physiology, Johns Hopkins University School of Medicine, Baltimore, MD 21205, USA; Solomon H. Snyder Department of Neuroscience, Johns Hopkins University School of Medicine, Baltimore, MD 21205, USA; Structural Genomics Consortium, University of Toronto, Toronto, ON M5G 1L7, Canada; Princess Margaret Cancer Centre and Department of Medical Biophysics, University of Toronto, Toronto, ON M5G 1L7, Canada

**Keywords:** Huntington disease, huntingtin, poly ADP-ribose, DNA repair, neurodegenerative disease, human, cerebrospinal fluid

## Abstract

Huntington disease (HD) is a genetic neurodegenerative disease caused by CAG expansion in the *Huntingtin (HTT)* gene, translating to an expanded polyglutamine tract in the huntingtin (HTT) protein. Age at disease onset correlates to CAG repeat length but varies by decades between individuals with identical repeat lengths. Genome-wide association studies link HD modification to DNA repair and mitochondrial health pathways. Clinical studies show elevated DNA damage in HD, even at the premanifest stage. A major DNA repair node influencing neurodegenerative disease is the PARP pathway. Accumulation of poly ADP-ribose (PAR) has been implicated in Alzheimer and Parkinson diseases, as well as cerebellar ataxia. We report that HD mutation carriers have lower cerebrospinal fluid PAR levels than healthy controls, starting at the premanifest stage. Human HD iPSC-derived neurons and patient-derived fibroblasts have diminished PAR response in the context of elevated DNA damage. We have defined a PAR-binding motif in huntingtin, detected huntingtin complexed with PARylated proteins in human cells during stress, and localized huntingtin to mitotic chromosomes upon inhibition of PAR degradation. Direct huntingtin PAR binding was measured by fluorescence polarization and visualized by atomic force microscopy at the single molecule level. While wild type and mutant huntingtin did not differ in their PAR binding ability, purified wild type huntingtin protein increased *in vitro* PARP1 activity while mutant huntingtin did not. These results provide insight into an early molecular mechanism of HD, suggesting possible targets for the design of early preventive therapies.

**Significance statement:** A consensus on dysfunctional DNA repair has emerged in neurodegenerative disease research, with elevated poly ADP-ribose (PAR) signaling more recently implicated. In contrast, we have identified a deficient PAR response in Huntington’s disease (HD) patient spinal fluid samples and cells. This may be explained by the inability of huntingtin protein bearing the HD-causing mutation to stimulate production of PAR the way the wild type protein does. Since drugs that target PAR production and degradation have already been developed, these findings present an exciting avenue for therapeutic intervention for HD.

## Introduction

Huntington disease (HD) is an autosomal dominant genetic neurodegenerative disease caused by a CAG expansion in exon1 of the *Huntingtin (HTT)* gene, which translates to an expanded polyglutamine tract in the huntingtin (HTT) protein. HD is characterized by psychiatric, cognitive, and motor disturbances for which some symptom management treatments, but no disease-modifying therapies, exist. Age at disease onset is negatively correlated with CAG expansion length, and onset typically occurs in the third or fourth decades of life [1]. However, age at disease onset in individuals with the same CAG length can vary by decades [2,3]. This variability has an inherited component [4,5], suggesting that other genes may act as modifiers of disease onset. Genome-wide association studies (GWAS) have primarily implicated DNA repair and maintenance pathways as such modifiers [2,3,6,7]. While the pathogenic mechanisms have yet to be elucidated, single nucleotide polymorphisms in DNA repair genes that affect somatic instability of the huntingtin locus have been linked to age at onset [8–10], and elevated levels of DNA damage in HD patient-derived samples have been reported [11–22].

The connection between DNA repair and neurodegeneration is not unique to Huntington disease, and has been described for Alzheimer disease [23], Parkinson disease [24], spinocerebellar ataxias [25,26], and amyotrophic lateral sclerosis [27,28]. Inherited mutations in a number of DNA repair genes cause neurodegenerative disorders [29–33], and of the nine neurodegenerative genetic polyglutamine diseases, seven of the causative proteins have been implicated in the DNA damage response [34]. Huntingtin forms a transcription-coupled DNA repair complex with RNA polymerase II subunit A, polynucleotide kinase-phosphatase (PNKP), and ataxin-3, and mutant huntingtin interferes with this function [21,22]. We have previously reported that huntingtin interacts with ataxia-telangiectasia mutated (ATM), and that ATM kinase activity is required for huntingtin-chromatin interaction in response to oxidative stress [12]. We have more recently identified a similar relationship between ATM and ataxin-1, the protein product of the *ATXN1* gene mutated in spinocerebellar ataxia 1 [35]. ATM signaling is also dysregulated in HD models and brain tissue [11], placing it as a DNA damage repair node in neurodegenerative diseases.

Another such node of DNA damage repair is PARP signaling, which has been implicated in Alzheimer, Parkinson, and Huntington diseases (reviewed in [36]) as well as progressive cerebellar ataxia [32] and amyotrophic lateral sclerosis (ALS) [28]. Of the 17 members of the PARP family, PARP1 and PARP2 are critical in the DNA damage response, with PARP1 accounting for 80-90% of DNA repair-related activity [37,38]. Upon activation by DNA breaks, PARP1 and PARP2 use ADP-ribose units from nicotinamide adenine dinucleotide (NAD^+^) as building blocks to generate poly ADP-ribose (PAR) chains of varying length and branching structure. PAR chains act as recruitment scaffolds for DNA repair proteins [39–41]. Part of the repair process involves PAR degradation by poly ADP-ribose glycohydrolase (PARG), which is required to allow DNA repair proteins access to the damaged DNA [42], and to recycle ATP precursor metabolites [43,44].

Excessive PAR polymerization, or lack of degradation, results in NAD^+^ depletion, energy crisis, and cell death by necrosis [45]. Overactivation of PARP1 can also trigger the non-apoptotic programmed cell death termed parthanatos [46], which is particularly important in neurodegenerative disease [47]. Hyper-PARylation has been implicated in several neurodegenerative diseases [32,48–50], including HD [51–53].

Here, we show that PAR signaling is dysregulated in HD patient cells and cerebrospinal fluid (CSF). Analysis of HD mutation carrier CSF samples revealed lower total PAR levels than control samples, a difference evident from the premanifest stage. Elevated DNA damage levels in HD iPSC-derived neurons and patient-derived fibroblasts were not reflected by increased PAR levels, and HD patient-derived cells had lower PARP1/2 inhibitor IC50 than control cells. Consistent with this, wild type purified recombinant huntingtin increased autoPARylation of purified recombinant PARP1 in an *in vitro* assay, while mutant huntingtin was deficient in this capacity. We detected endogenous huntingtin complexed with PARylated proteins in human retinal epithelial cells, which prompted analysis of the huntingtin structure and identification of a PAR-binding motif. Fluorescence polarization and atomic force microscopy revealed a direct interaction between huntingtin protein and PAR polymers. While PAR binding by wild type or mutant huntingtin did not differ, mutant huntingtin was not able to stimulate PARP1 activity while wild type protein demonstrated this activity. This provides a possible mechanistic link to the observations made in patient cells and CSF. In addition to uncovering a role for the normal huntingtin protein in PAR biology, these results provide insight into the very early molecular mechanism of HD pathogenesis, to potentially reveal targets for an early preventive therapy for HD.

## Results

### The PAR response is deficient in HD patient-derived samples

Elevated PAR levels have been observed in human samples from multiple neurodegenerative diseases [32,48–50]. Although PARP1 inhibition was reported to be neuroprotective in a mouse model of HD [51,52], and elevated PARP1 levels were observed in the neurons and glia of the caudate nucleus from HD patients [53], it remains unknown whether hyper-PARylation contributes to HD pathology. As cancer drugs targeting PARP1 are under consideration for repurposing to treat neurodegenerative diseases [54,55], investigation of their potential for HD is imperative. To test patient-relevant CNS-specific samples, we measured PAR levels in CSF from HD patients compared to controls. Samples were obtained from the HD-CSF study, an 80-participant cross-sectional study of HD mutation carriers and matched healthy controls [56,57] (see methods section for details). In contrast to similar analyses in Parkinson’s disease [50], we found that PAR levels were significantly lower in the CSF from both pre-manifest (p = 0.0001) and manifest HD patients (p = 0.0004) compared to controls, and that gene status had a large effect on PAR levels in CSF (η^2^ = 0.35) (**Fig 1**). PAR levels in the CSF of HD mutation carriers did not correlate with CSF mutant huntingtin or neurofilament light chain (NfL) levels, both of which were shown to correlate with disease severity in HD patients [56,57] (**Fig S1**). In addition, CSF PAR levels in HD mutation carriers did not correlate with clinical measures of disease progression such as disease burden score, total functional capacity, or total motor score (**Fig S2**). Yet, lower PAR levels correlated with the presence of the mutant huntingtin allele.

**Figure 1:**
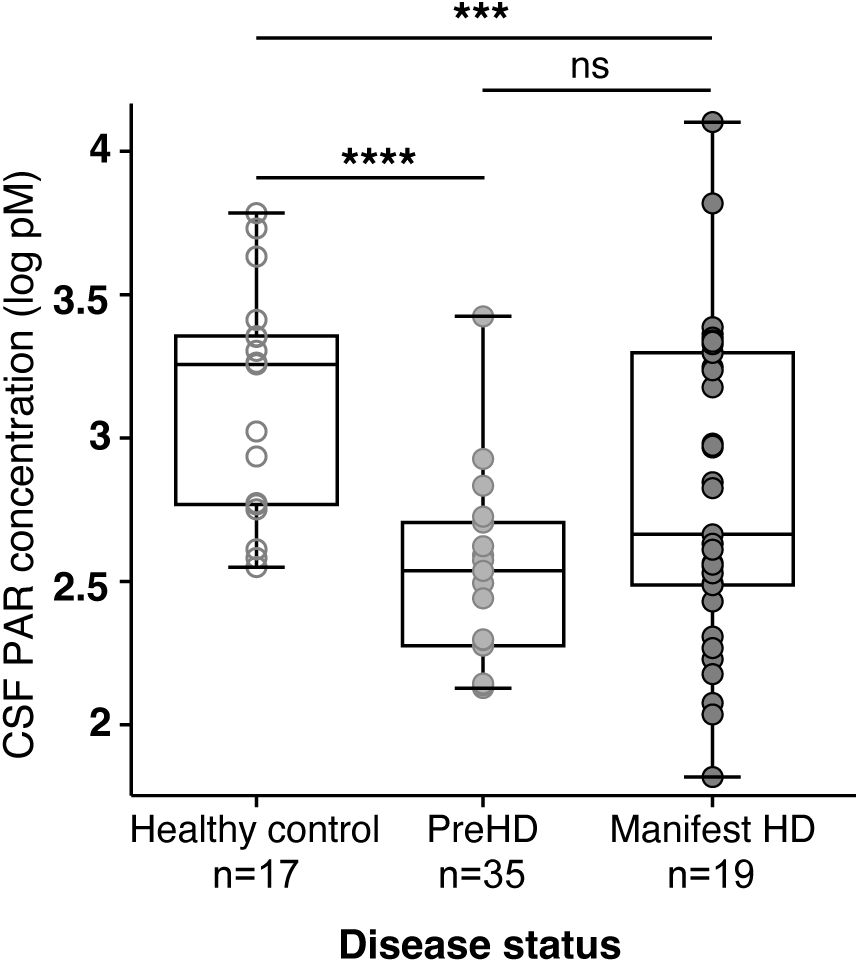
PAR levels are reduced in premanifest and manifest HD patient CSF. CSF samples from control, premanifest HD, and manifest HD subjects were blinded and analyzed for PAR levels by ELISA. Group comparisons were assessed using multiple regression with post-hoc Wald tests. *Survives Bonferroni correction for multiple comparisons.

The reduced PAR levels in the CSF of premanifest and manifest HD patients were unexpected given the numerous reports of increased DNA damage in HD patient samples and models. For example, iPSC-derived neurons bearing the HD mutation have increased DNA breaks and expression of damage markers phospho-ATM, _γ_H2AX, and phospho-p53 [21,22]. We therefore aimed to determine whether this increase in DNA damage was reflected by elevated PAR levels. As shown in **Fig 2A**, we found no corresponding increase in PAR levels in HD iPSC-derived neurons. In fact, neurons bearing a juvenile onset HD allele (Q77) had lower PAR levels than control, consistent with the observations made in patient CSF.

**Figure 2:**
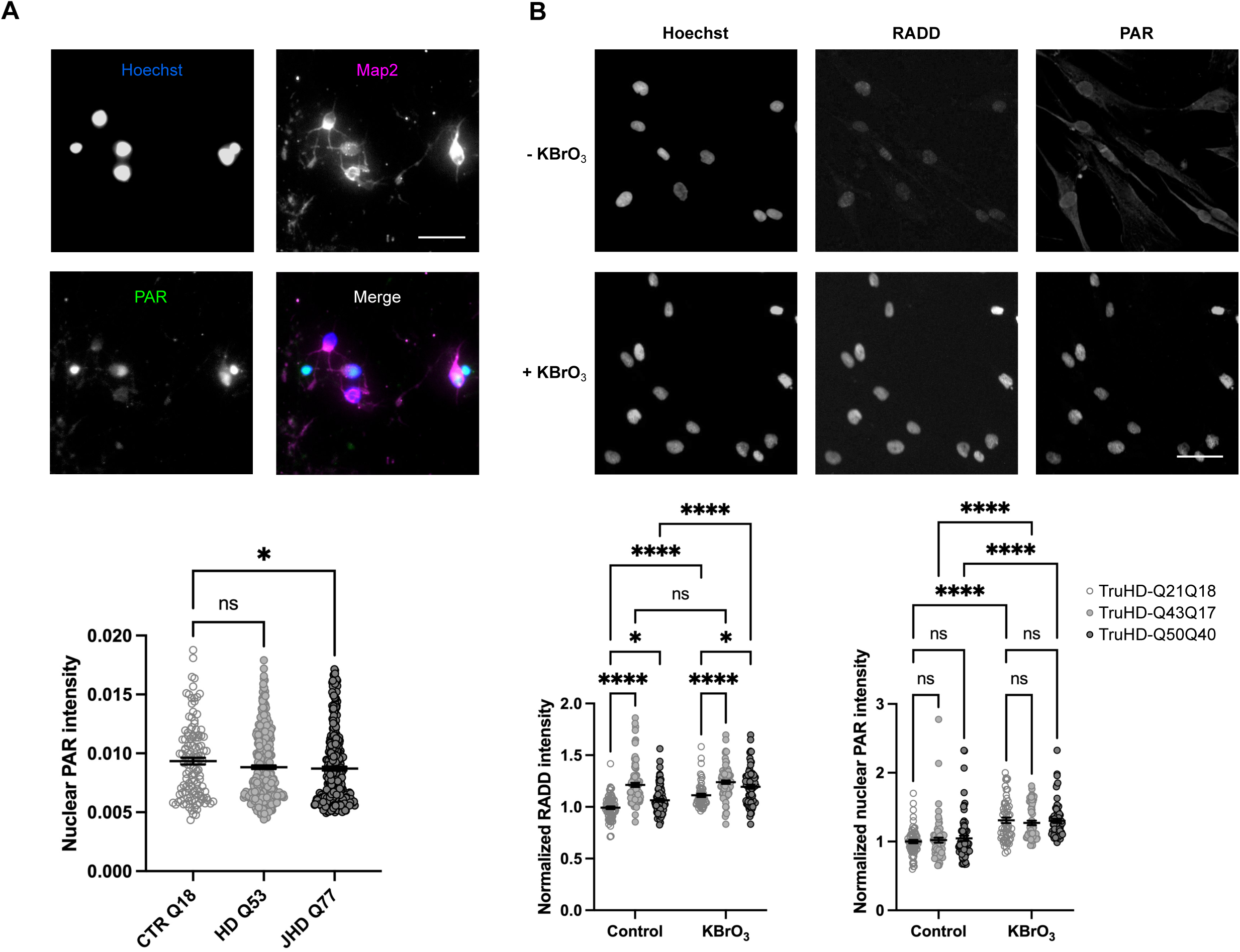
The PAR response is deficient in HD cells. **>A**: iPSC-derived neurons were fixed and stained with neuronal marker Map2 (red) and MABE1031 PAR detection reagent (green). Nuclear PAR intensity in Map2-positive cells was measured using CellProfiler. Data from six (CTR Q18 and HD Q53) or four (JHD Q77) differentiation replicates are shown (n=100-300 nuclei per cell line). Results were analyzed by Kruskal-Wallis test and corrected for multiple comparisons using Dunn’s test. Error bars: SEM. **B**: hTERT-immortalized fibroblasts from healthy control (TruHD- Q21Q18) and HD patients (TruHD-Q43Q17 and TruHD-Q50Q40) were treated with 100 mM KBrO_3_ for 30 min followed by Repair Assisted Damage Detection (RADD) to detect DNA damage, and co-staining with MABE1031 PAR detection reagent. Representative images of TruHD-Q21Q18 cells are shown. Nuclear RADD and PAR intensity were measured using CellProfiler, mean intensity recorded for each image (18 images per condition; >500 cells), and values normalized to the control condition. Data from three independent experiments is shown. Results were analyzed by two-way ANOVA and corrected for multiple comparisons using Tukey’s test. Error bars: SEM. Scale bars: 50 microns.

We and others have previously reported elevated levels of DNA damage in fibroblasts derived from HD patients compared to controls [12,18,20]. We therefore measured DNA damage and PAR levels in control cells bearing wild type alleles that encode 21 or 18 polyglutamine residues (TruHD-Q21Q18), and HD patient-derived cells carrying one expanded allele (TruHD-Q43Q17), or two expanded alleles (TruHD-Q50Q40), TruHD cells [58]. We employed Repair Assisted Damage Detection (RADD) [59] to measure DNA breaks and PAR levels in the same cells. RADD directly detects DNA damage using DNA processing enzymes to detect and modify sites of DNA damage for a subsequent gap filling fluorescent labeling reaction. Like HD iPSC-derived neurons, hTERT-immortalized HD patient-derived fibroblasts did not exhibit the increase in PAR levels expected to occur in the context of elevated DNA damage (**Fig 2B**). This was the case under conditions of oxidative stress, which we have previously shown to be associated with elevated DNA damage in these cells by comet assay [12], as well as under basal conditions, in which increased damage was also detected by RADD (Fig 2B) .

To ensure that the PAR detection assay is within the dynamic range and the signal has not reached saturation, we quantified PAR signal intensity in a KBrO_3_ dose response experiment. As shown in Fig S3, a treatment of 100 mM KBrO_3_ for 30 minutes produces a PAR response within the dynamic range. Further, HD cells exhibited lower PAR levels than wild type cells at the other KBrO_3_ doses.

Since hTERT expression is equal across the immortalized fibroblast lines [58], and similar results were obtained in iPSC-derived neurons and patient CSF, we did not anticipate a confounding effect of hTERT overexpression. This was confirmed by repeating the experiment in the primary fibroblasts from which the TruHD lines were derived (**Fig S4**). It should be noted that although we did not observe the expected elevated PAR phenotype in HD patient-derived cells, PAR was generated in response to oxidative stress, indicating that they are not fully deficient in this capacity (**Fig 2B**). To further characterize this PAR signaling deficiency, we correlated per-nucleus RADD and PAR signal intensities for control and HD patient-derived cells and found them to be correlated (**Fig S5**). As expected, the correlation increased with KBrO_3_ treatment, however HD cells had lower Spearman r values than wild type cells under both conditions. This suggests that the PAR response is not fully deficient but is subdued in the HD context, as would be expected for a late onset, slowly progressing disease. Together, the reduced PAR levels in HD patient CSF, and lack of elevated PAR in the context of elevated DNA damage in HD neurons and patient-derived fibroblasts, suggest that the PAR response is deficient in HD patients.

We next examined the mechanisms of PAR production and degradation in TruHD cells. During the DNA repair process, PAR is rapidly generated by PARP1 and PARP2, and degraded by enzymes such as PARG. To determine whether the lower than expected PAR levels were due to increased PARG activity, we performed dose response experiments to measure the PARG inhibitor PDD00017273 IC50 [60] and found no difference between HD and control cells (**Fig 3A**). In contrast, dose response experiments with the PARP1/2 inhibitor veliparib revealed lower veliparib IC50 in HD cells compared to controls (TruHD-Q21Q18 IC50 = 140 nM; TruHD- Q43Q17 IC50 = 60 nM; TruHD-Q50Q40 IC50 = 60 nM) (**Fig 3B**). This is despite similar levels of PARP1, PARP2, and PARG across cell lines (**Fig S6A**) and equal PARP1 chromatin retention upon oxidative stress (**Fig S6B**). To ensure that the lowered IC50 is not an artifact specific to veliparib, we repeated the experiment with talazoparib, which acts by a different mechanism than veliparib [61,62], and found a reduced IC50 in HD cells (Fig S6C), as was seen with veliparib. To address whether the reduced PARP inhibitor IC50 values were a direct effect of huntingtin, we tested the autoPARylation activity of purified recombinant PARP1 in the presence of purified recombinant HTT-HAP40 Q23 or HTT-HAP40 Q54. As shown in **Fig 3C**, wild type huntingtin stimulated PARP1 activity in a dose-dependent manner, while mutant huntingtin had no effect. This provides a possible mechanistic link to the reduced PARP inhibitor IC50 values, and subdued PAR response observed in HD patient-derived fibroblasts, as well as the reduced PAR levels in HD iPSC-derived neurons and CSF from HD patients.

**Figure 3:**
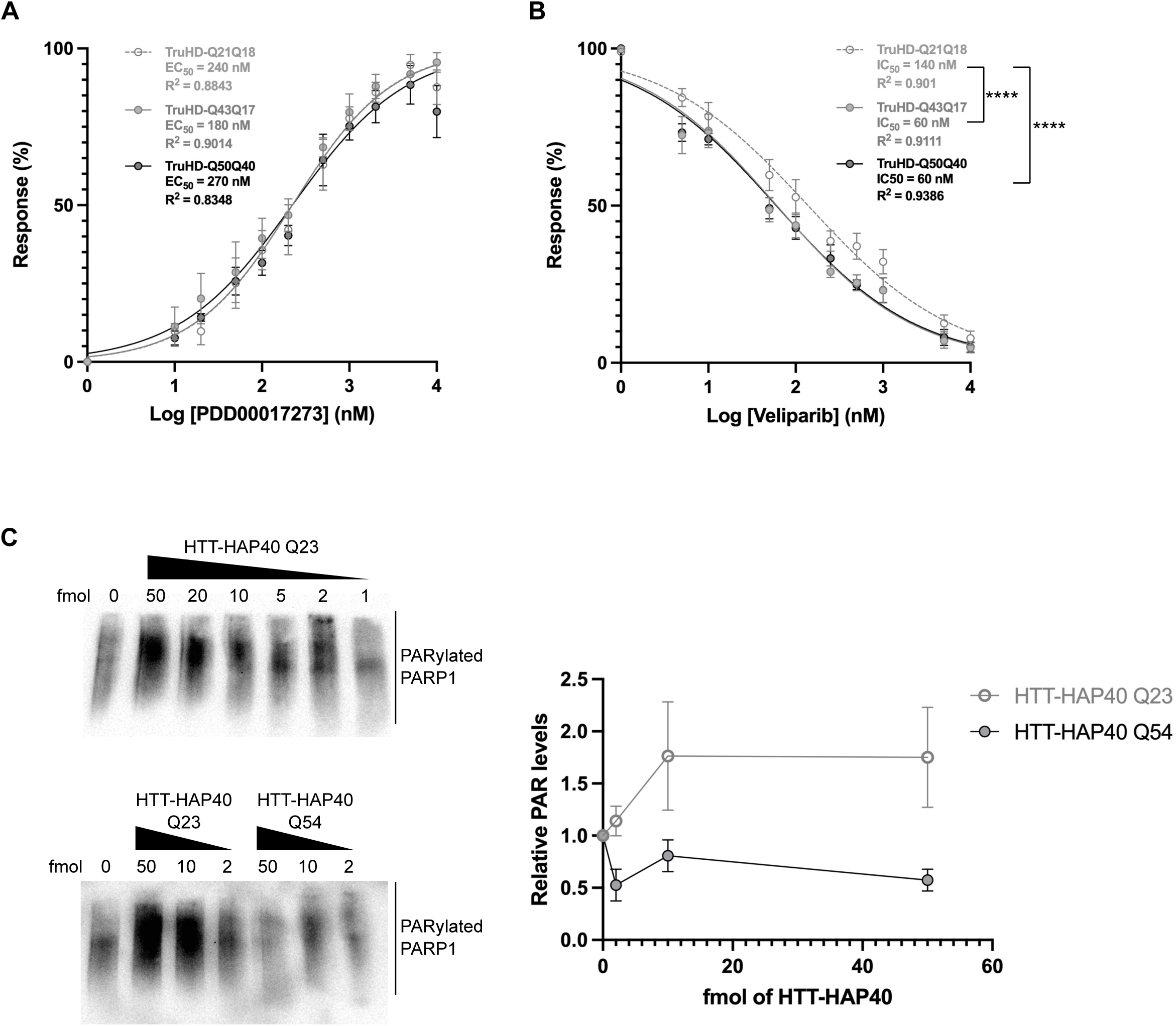
PARP1/2 activity is higher in the presence of wild type huntingtin. Fibroblasts were pre-treated with increasing doses of PDD00017273 (**A**) or veliparib (**B**) for 30 min followed by 100 mM KBrO_3_ for 30 min in the presence of inhibitor. Veliparib dose response was carried out in the presence of 5 _μ_M PARG inhibitor to enable pan-ADP-ribose detection by MABE1016. EC_50_ and IC_50_ values were calculated from nuclear PAR staining intensity (10-12 images per condition; >800 cells) using GraphPad Prism. Error bars = SEM for 4 (PARG inhibitor) or 8 (veliparib) experiments. **** p<0.0001 (Brown-Forsythe and Welch ANOVA tests). **C**: 10 fmol recombinant PARP1 was incubated with the indicated amounts of recombinant HTT-HAP40 for 2 hours at 30°C. Reactions were separated by SDS-PAGE and immunoblotted with MABE1016 pan-ADP-ribose detection reagent. Signal intensities were quantified using ImageJ. Values from three (HTT-HAP40 Q54) or four (HTT-HAP40 Q23) experiments are shown.

### Huntingtin interacts with PARylated proteins

Mutations associated with progressive cerebellar ataxia with oculomotor apraxia type 1 (AOA1) result in reduced expression of the scaffolding protein XRCC1, causing persistent unrepaired DNA damage and concomitant prolonged PARP1 activity [32]. Since PAR binding is important for the scaffolding function of XRCC1 in DNA repair complex formation [63], we sought to determine whether huntingtin could also bind PAR.

We first interrogated the results of a mass spectrometry study identifying proteins that interact with huntingtin under conditions of oxidative stress (Table S1). The list of huntingtin interactors was compared to a compiled list of PARylated proteins from three independently generated datasets (**Fig S7**) [64–66]. As shown in **Figure 4**, 122 of the 298 (41%) huntingtin-interacting proteins were also found in a database of PARylated proteins. Fisher’s test returned a significance of 2.1 x 10^-81^. Thus, a significant proportion of huntingtin-interacting proteins are reported to be modified by PAR (Table S2).

**Figure 4:**
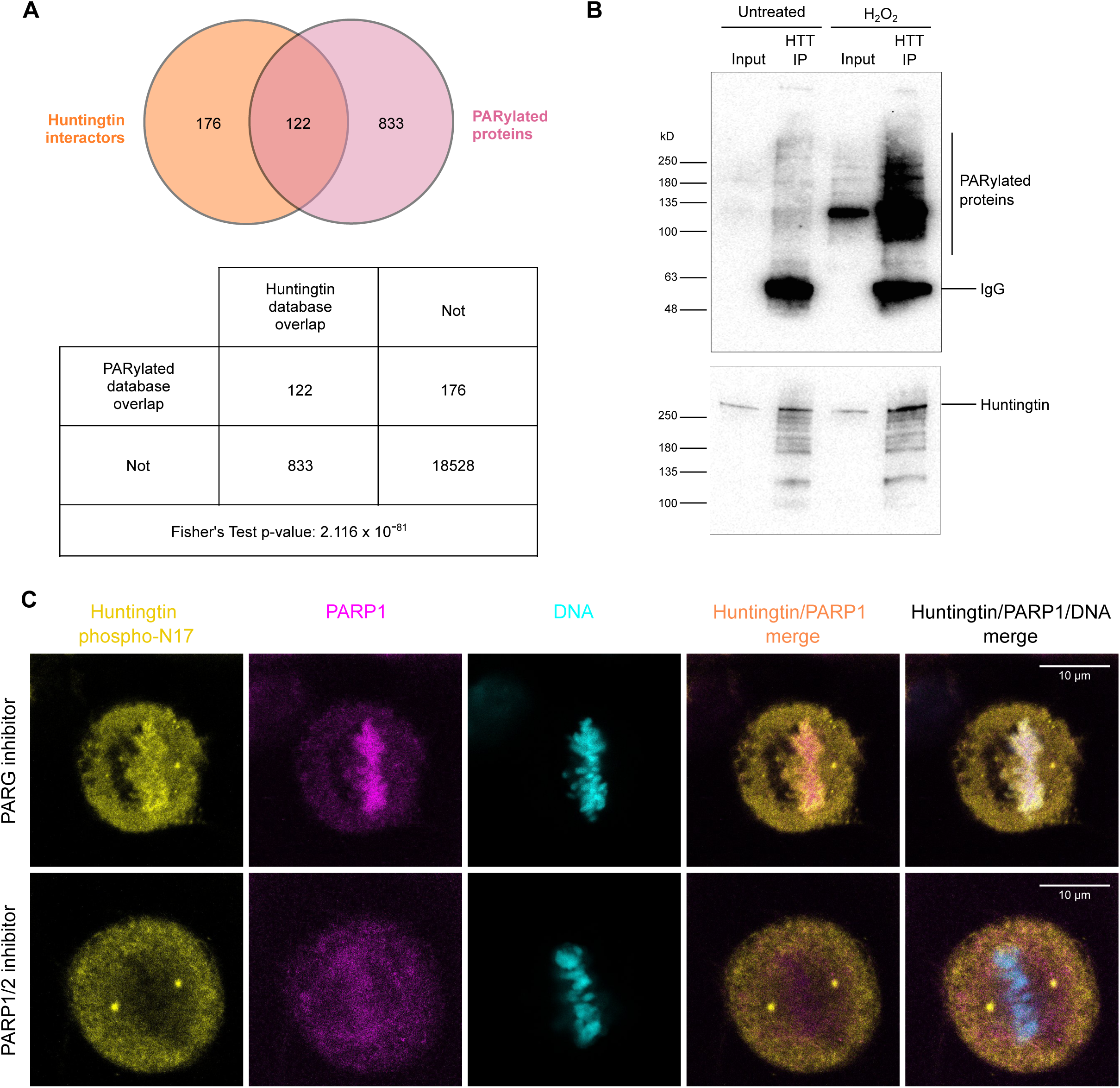
Huntingtin interacts with PARylated proteins. **A**: Degree of overlap between huntingtin interacting proteins and a list of PARylated proteins compiled from three independent studies, with Fisher’s exact test for statistical significance. **B**: RPE1 cells were treated with 400 _μ_M H_2_O_2_ in HBSS for 10 min and proteins crosslinked with 1% PFA prior to lysis. Huntingtin was immunoprecipitated with EPR5526 and associated proteins separated by SDS-PAGE and immunoblotted with the indicated antibodies. PARylated proteins of various sizes in the whole cell lysate (input) and anti-huntingtin immunoprecipitate (HTT IP) were detected with pan ADP-ribose detection reagent (MABE1016) followed by HRP-conjugated anti-rabbit secondary antibody. Rabbit IgG signal from the anti-huntingtin immunoprecipitating antibody (EPR5526) was visible upon incubation with secondary anti-rabbit antibody. Results representative of four experiments. **C**: RPE1 cells were treated with either 10 _μ_M PDD00017273 PARG inhibitor (top panel) or 1 _μ_M talazoparib PARP1/2 inhibitor (bottom panel) for 40 min prior to methanol fixation and immunofluorescence against huntingtin phosphorylated at residues S13 and S16 within the N17 domain (Huntingtin phospho-N17, yellow), and PARP1 (PARP1, magenta), followed by counterstaining with Hoechst (DNA, cyan). Image representative of all mitotic cells observed (n > 10 cells from two independent experiments). Scale bar: 10 _μ_m.

We next asked whether huntingtin exists in complex with PARylated proteins in cells. Since fibroblasts do not produce sufficient protein for immunoprecipitation analyses, we turned to RPE1 cells, which are hTERT-immortalized and therefore retain DNA repair pathway function. As expected, PAR levels were increased in RPE1 cells treated with hydrogen peroxide (**Fig 4B**). Furthermore, PARylated proteins were enriched in huntingtin immunoprecipitates upon oxidative stress as shown by western blot analysis. This is consistent with the high degree of overlap between datasets of huntingtin-interacting proteins and PARylated proteins.

To further investigate the relationship between huntingtin and PAR in human cells, we examined the subcellular localization of endogenous huntingtin upon manipulation of PAR production and degradation. While endogenous huntingtin localization in interphase cells was similar across conditions, we found that in mitotic cells treated with PARG inhibitor, huntingtin phosphorylated at residues S13 and S16 localized strongly to condensed mitotic chromosomes (**Fig S8** and **Supplementary video 1**). This staining pattern is strikingly similar to that of PAR itself during mitosis [67,68], and provides further evidence that huntingtin binds PAR in cells. In contrast, upon inhibition of PARP1/2 activity, huntingtin localized primarily to the mitotic spindle poles (**Fig 4C**), as we and others have seen previously in untreated cells [69,70]. Thus, detection of huntingtin-PAR complexes by immunoprecipitation in cell lysates, and by immunofluorescence in intact cells, suggests that normal huntingtin may play a role in PAR biology, and raises the possibility that huntingtin may directly bind PAR.

### Huntingtin contains a PAR-binding motif and directly binds PAR

We examined the huntingtin sequence for potential PAR-binding motifs (PBMs) according to consensus sequences derived by others [64,71] and found five putative PBMs (**Fig 5A**). While huntingtin putative PBMs did not match the consensus with 100% similarity, their consensus fitting was comparable to those from previously validated PAR-binding proteins including DNA dependent protein kinase catalytic subunit (DNA-PK) [71] and 60 kD SS-A/Ro ribonucleoprotein [72].

**Figure 5:**
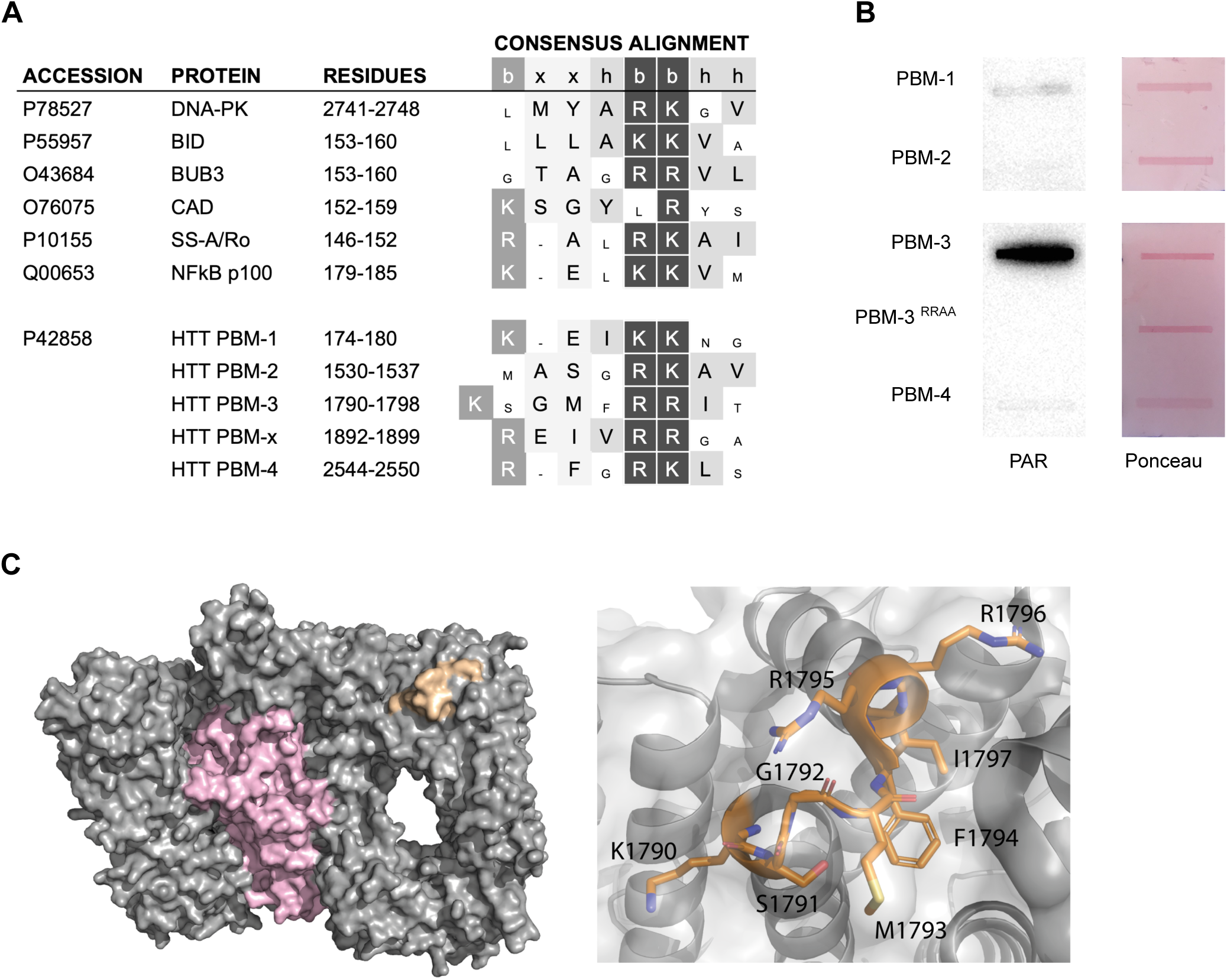
Huntingtin has a PAR-binding motif (PBM) **A**: Known PBMs compared to putative PBMs in huntingtin. PBM-X is not solvent-accessible and was not analyzed further. b: basic, h: hydrophobic, x: any amino acid. Critical basic amino acids depicted in black boxes. **B**: Peptides were slot-blotted onto nitrocellulose then overlaid with 0.2 _μ_M PAR polymer. After washing, anti-PAR western was performed with pan ADP-ribose detection reagent MABE1016. **C**: Left: high resolution cryoEM model of huntingtin-HAP40 complex (PDB - 6X9O) shown in surface representation with huntingtin in gray, HAP40 in pink, and the PAR-binding motif in orange. Right: PAR-binding motif shown in stick representation in orange. Positively charged K1790, R1795 and R1796 residues are surface exposed.

To test these putative motifs, we performed PAR overlay assays with peptides representing the putative PBMs (**Fig 5B**). PBM-3 displayed strong PAR-binding activity, which was ablated by mutation of the critical arginine residues. Multiple sequence alignment of human huntingtin and several orthologous species revealed a high degree of evolutionary conservation of PBM-3 and the surrounding sequences (**Fig S9**). Mapping of this sequence on the cryo-EM structure of huntingtin [73] shows that PBM-3 is solvent accessible (**Fig 5C**), situated on the bridge domain at the interface of the N-HEAT domain with a surface area of ∼890 Å^2^. The motif spans a connecting loop region in this HEAT repeat in addition to a small section of each of the two flanking α-helices. The positively charged K1790, R1795 and R1796 residues are surface exposed in this model of the structure, indicating how they might interact with negatively charged PAR molecules.

We then asked whether the huntingtin PBM-3 bears resemblance to the PBM from the structurally similar [74] and functionally related [12] protein, ATM. Similar to ATM, the PAR-binding motif of huntingtin is a surface-exposed helix-turn-helix within a HEAT repeat (**Fig S10**). These results suggest that huntingtin PBM-3 is a *bona fide* PAR-binding motif, and that at least some of the interactions detected by immunoprecipitation and mass spectrometry (**Fig 4** and Table 1) could be direct.

To determine whether huntingtin directly interacts with PAR, we tested purified full-length huntingtin protein and PAR linear polymer in a fluorescence polarization assay. As shown in **Fig 6A**, huntingtin binds long (26-mer) PAR chains, in preference to shorter (11-mer) substrates.

**Figure 6:**
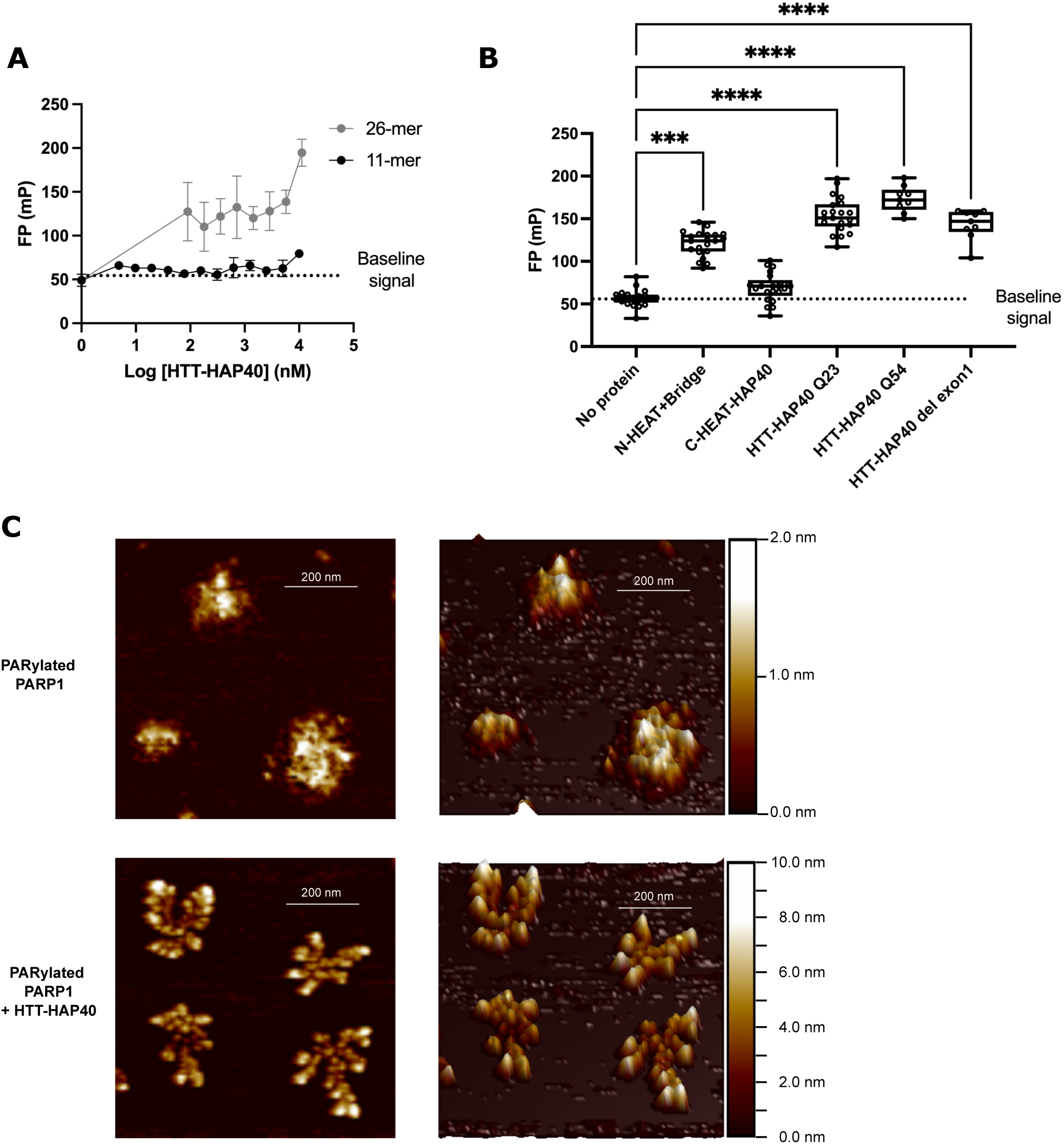
Huntingtin directly binds PAR. **A**: Fluorescence polarization assays using FAM-labeled 11-mer or 26-mer PAR and purified HTT-HAP40 Q23. Results for 2 experiments are shown. Error bars = SD. **B**: Fluorescence polarization assays using FAM-labeled 26-mer PAR and 10 _μ_M of the indicated subdomains of huntingtin. Reactions were carried out with 3-4 intra-assay replicates. Results for 3 experiments were analyzed by one-way ANOVA and corrected for multiple comparisons by Kruskal-Wallis test (***p<0.0005, ****p<0.0001). **C**: Recombinant HTT-HAP40 was added to PARP1 activity assays and reactions were deposited on mica and visualized by atomic force microscopy. 2D (left) and 3D (right) images are shown with corresponding colour scale for PARylated PARP1 (2.0 nm) or PARylated PARP1 with HTT-HAP40 (10.0 nm).

This is similar to the substrate length specificity of other PAR binding proteins [75]. It should be noted that while these results support a direct interaction, the technical constraints of the experiment preclude saturation of the binding curve and therefore determination of binding kinetics and specificity. At 10 μM protein, HTT-HAP40 Q23 exhibited lower binding than the positive control PAR-binding protein HUWE1, but higher binding than the negative control non-PAR-binding protein USP5 (**Fig S11**). We next tested the PAR binding ability of different huntingtin subdomain constructs [76], and found that a construct comprising the N-HEAT and Bridge domains, which contains PBM-3, binds 26-mer PAR, while the C-HEAT domain (in complex with HAP40) does not (**Fig 6B**). We did not detect a significant difference in PAR binding between wild type and mutant huntingtin, nor upon deletion of the exon 1 domain in this assay (**Fig 6B**). Thus, a difference in PAR binding does not explain the differential effect that wild type and mutant huntingtin have on PARP1 activity *in vitro* (**Fig 3C**). These results suggest that huntingtin can directly bind linear PAR chains of at least 26 ADP-ribose units in length, and that binding may be mediated through PBM-3.

In order to test whether huntingtin may directly bind PAR chains of various sizes and branching structures, we included purified huntingtin protein in *in vitro* PARP1 reactions, and visualized the reactions by atomic force microscopy at the single molecule level. In the absence of huntingtin, we observed auto-PARylated PARP1 structures consistent with previous reports [77–79] (**Fig 6C**). In contrast, the presence of huntingtin protein in the PARP1 reactions resulted in large rosette structures consistent with huntingtin protein bound to PAR chains (**Fig 6C**).

While visualization of the PARP1 reaction in the presence of huntingtin may give insight into the increased PARP1 autoPARylation activity we detected in the presence of wild type huntingtin by western (**Fig 3C**), differing buffer conditions and mica substrate effects preclude the elucidation of precise mechanism, which is the topic of future studies.

Taken together, these results indicate that huntingtin binds directly to PAR chains *in vitro*, which accounts for its interaction with PARylated proteins in cells. This establishes a link between huntingtin and PAR signaling biology. Despite equivalent PAR binding, wild type huntingtin increases *in vitro* PARP1 activity while mutant huntingtin does not. This may explain the reduced PAR levels in HD patient CSF, and the subdued PAR response and reduced PARP inhibitor IC50 values we observed in patient-derived cells.

## Discussion

The huntingtin protein is a large scaffold that participates in numerous cellular processes [80]. We have previously defined a role for huntingtin in the response to oxidative DNA damage [12], and characterized the nuclear/primary cilium localization signal [81] and nuclear export signals [82,83] regulating its translocation from the endoplasmic reticulum, where it is tethered by the N17 domain, to the nucleus [69,84,85]. Here, we define an evolutionarily conserved PAR-binding motif and show that huntingtin interacts with PARylated proteins. *In vitro*, wild type purified recombinant huntingtin increases autoPARylation of purified recombinant PARP1, while mutant huntingtin is deficient in this capacity. This may contribute to the reduced levels of PAR we observed in the CSF from HD patients and the deficient PAR levels seen in HD iPSC-derived neurons and HD patient-derived fibroblasts in the context of elevated DNA damage.

We have previously hypothesized [34,86] that aberrant mutant huntingtin function in DNA repair plays a role in the elevated levels of DNA damage seen in several HD models and tissues [11–22]. The ability of wild type but not mutant huntingtin to stimulate PARP1 autoPARylation activity provides one possible contributing mechanism. We identified and characterized a PAR binding motif within huntingtin, and found that mutant huntingtin PAR binding was not different from that of wild type. This indicates that mutant huntingtin can bind PAR but cannot stimulate its activity the way that wild type huntingtin can. . While the identification of PBM-3 as a structurally and evolutionarily conserved PAR binding motif strongly suggests that it plays a role in huntingtin PAR binding, the data presented do not exclude the possibility that additional contact points contribute to PAR binding, including possibly PBM-1, which is also found in the N-HEAT subdomain and showed weak binding in the PAR overlay assay. We show that huntingtin directly binds PAR through the identification of a defined PAR binding motif, fluorescence polarization and atomic force microscopy, as well as an interactome of over 100 PARylated proteins and huntingtin localization to PAR-coated mitotic chromosomes upon PARG inhibition.

Dysregulation of PAR signaling has now been linked to multiple neurodegenerative diseases. A common theme among these disorders is the hyperactivation of PARP1 in response to oxidative damage. This may also be the case in HD, as evidenced by strong PARP1 expression in the neurons and glia of caudate nucleus from affected HD patients [53], and by the beneficial effect of PARP1 inhibition in the R6/2 mouse model [51,52]. In contrast, we detected lower PAR levels in HD patient CSF and reduced PARP1/2 inhibitor IC50 in fibroblasts from HD patients. The decreased PAR levels we detected in the CSF from HD patients, an anomaly by comparison to similar studies in different neurodegenerative diseases [32,48–50], nonetheless reflect dysregulated PAR signaling in HD. This may parallel the paradoxical PARP inhibitor and PARG inhibitor cancer treatment options [87], which show that tipping the balance in either direction is detrimental to cancer cells. Static measurements of hypo- and hyper-PARylation may not reflect the dynamic nature of PAR metabolism, but may instead indicate PAR signaling dysregulation in HD. While CSF PAR levels did not correlate with disease measures, the robust reduction in PAR, even in premanifest patients, provides clinically relevant evidence for a role of PAR signaling dysregulation in the early stages of disease.

Although we also observed lower than expected PAR levels in iPSC-derived neurons, the important caveat is that CSF PAR is not intraneuronal PAR. The clinical data from CSF may only indicate that there is an abnormal PAR dynamic process, hence an impaired ability to repair DNA, as seen clinically by others as early as premanifest HD [13,14].

It is interesting to speculate on why PAR levels in the CSF are lower in HD mutation carriers while there is no difference in HD patient-derived fibroblast intracellular levels. This could reflect differences between systemic nervous system fluids versus skin-derived cells grown in a dish, whereby cell culture conditions require a minimum PAR level for cell proliferation. Basal PAR levels were in fact lower in iPSC-derived cultured, non-dividing neurons representing the more severe juvenile HD case (Q77), consistent with lower PAR levels in CSF from HD patients. The ELISA method used to measure CSF PAR levels recognizes both PARylated proteins and free PAR released by brain cells during waste clearance.

The fact that PAR levels did not correlate with biomarker levels or clinical measures of disease likely reflects the transient and fluctuating nature of PAR, meaning that it may not be useful as a biomarker of disease state. This is also true of other disease-related molecules such as inflammatory markers [88] that are linked to pathology but do not have optimal kinetics for biomonitoring. Further, where PAR may be relevant to regionally specific pathology, its level in CSF is not specific to a particular brain region. Therefore, any regional specific changes could be diluted by more general fluctuations. We speculate that dysregulation of PAR signaling in HD may have beneficial secondary consequences, such as the reduced cancer rates seen in CAG expansion carriers [89], which could possibly be related to reduced PARP1/2 activity. This could in turn contribute to the apparent evolutionary advantage of longer CAG repeat lengths, which are not in equilibrium in human populations, but are subject to mutational bias towards expansion [90].

An interaction between HD and cancer via PARP1/2 activity may provide opportunity for the repurposing of currently available cancer drugs targeting this pathway. Future studies will look at different classes of FDA-approved PARP1 inhibitor drugs as some trap PARP1 on DNA, and some can cross the blood brain barrier while others cannot [54,55]. Interrogation of large banks of HD patient data [91] may make it possible to determine changes in disease progression associated with administration of such drugs, while the emerging development of new PARP and PARG inhibitors may hold promise for HD and neurodegenerative diseases at large.

## Materials and Methods

### Antibodies and reagents

All reagents were from MilliporeSigma unless otherwise stated. Veliparib (ABT-888) was from Selleckchem. PARG inhibitor (PDD00017273) was from AdipoGen Lifesciences. Antibodies are listed in Supplementary methods. Mica and specimen supports were from Ted Pella Inc.

### Human CSF samples

CSF samples were collected as part of the HD-CSF study, a prospective single-site study with standardized longitudinal collection of CSF, blood and phenotypic data (online protocol: 10.5522/04/11828448.v1). The cohort included manifest and premanifest HD mutation carriers as well as healthy controls who were age and gender matched to the entire HD mutation carrier group (**Fig S12**). All samples were de-identified prior to use in this study. In summary, lumbar punctures were performed after 12-hour fasting, in the early morning, and samples were processed within 30 minutes after the collection. Processed samples were aliquoted and stored at -80°C until shipment on dry ice for PAR quantification.

### Measurement of CSF analytes

PAR quantification was performed using an in-house ELISA as previously described [50]. NfL, mHTT, and Hemoglobin were quantified using the Simoa® Neurology 4-plex B kits (Item 103345, Quanterix, Lexington MA, USA, at UCL), the SMCxPro (2B7/MW1, in-house at Evotec) and ELISA (E88-134, Bethyl Laboratories; at Evotec), respectively, as described previously (57).

### Cell culture and treatments

TruHD cells[58] were cultured in MEM (Life Technologies #10370) with 15% fetal bovine serum (FBS; Life Technologies) and 1X GlutaMAX (Life Technologies #35050) and grown at 37°C with 5% CO_2_ and 5-8% O_2_. ST*Hdh* cells[92] were cultured in DMEM (Life Technologies #11995) with 10% FBS and grown at 33°C with 5% CO_2_. RPE1 cells (American Type Culture Collection) were cultured in 1:1 DMEM/Nutrient Mixture F-12 (DMEM/F12; Life Technologies #11330) with 10% FBS and 0.01% hygromycin and grown at 37°C with 5% CO_2_ and 5-8% O_2_. Induction of DNA damage with H_2_O_2_ or KBrO_3_ was done in Hank’s balanced salt solution (HBSS) or phosphate-buffered saline (PBS), at the concentrations and durations indicated in figure legends.

### Neuronal differentiation of iPSCs

HD iPSCs were differentiated as previously described [21,93]. All samples were de-identified prior to use in this study.

### Measurement of PAR levels in cells

Cells were seeded in 8-well ibiTreat µ-Slides (Ibidi) to ∼95% confluence. For KBrO_3_ dose response experiments, cells were seeded in glass-bottom 96-well plates (CellVis). After the indicated treatments, cells were stained and imaged as described in Supplementary methods.

### Repair assisted damage detection (RADD)

Cells were seeded in 8-well ibiTreat µ-Slides (Ibidi) or Mod3D chambers [94] to ∼95% confluence. Cells were washed with PBS then incubated with PBS containing 0 mM or 100 mM KBrO_3_ for 30 min. RADD was performed as described [59] and Supplementary methods. Cells were imaged in PBS using the 20× objective on the EVOS FL Auto 2 widefield microscope.

Nuclei were identified as primary objects in CellProfiler [95] using the Hoechst staining, then pixel intensity of the RADD and PAR staining within nuclei was calculated and the mean intensity recorded for each image. Eighteen images per well were captured, representing >500 cells per experiment.

### Inhibitor dose response experiments

PARP1/2 and PARG activities were measured using the method described by James et al [60] and in the Supplementary methods. IC50 values were calculated using GraphPad Prism Version 9.4.1. Means were assessed by Brown-Forsythe and Welch ANOVA tests followed by Dunnett’s T3 multiple comparisons test with individual variances computed for each comparison.

### Immunofluorescence

To measure chromatin retention of PARP1, TruHD cells were grown in CellCarrier-96 Ultra Microplates (Perkin Elmer) and treated for 30 min with 100 mM KBrO_3_ dissolved in PBS containing Ca^2+^ and Mg^2+^. Soluble proteins were extracted with cold 0.2% Triton X-100 in PBS containing Ca^2+^ and Mg^2+^ for 2 min on ice prior to staining as described in Supplementary methods.

For imaging of mitotic cells, RPE1 cells were stained as described in Supplementary methods.

### Cell lysis and western analysis

For measurement of protein levels by western, cells were lysed using radioimmunoprecipitation assay buffer (50 mM Tris-HCl pH 8.0, 150 mM NaCl, 1% NP-40, 0.25% sodium deoxycholate, 1 mM EDTA) with protease and phosphatase inhibitors (Roche) and protein levels were measured with the Qubit Protein Broad Range (BR) Assay (ThermoFisher). Westerns were performed and quantified as described in Supplementary methods.

### Purification of huntingtin-interacting proteins

ST*Hdh*^Q7/Q7^ mouse striatal precursor cells were treated with either HBSS, HBSS containing 60 μg/mL methyl methanesulfonate (MMS) for 20 min, or HBSS containing 100 μM H_2_O_2_ for 1 hour. Proteins were purified as described in Supplementary methods.

### Mass spectrometry and protein identification

Samples were trypsin-digested, desalted on C18 column, and LC-MS/MS performed on the Q Exactive HF-X mass spectrometer (SPARC BioCentre, Toronto, Canada). For database searching, tandem mass spectra were extracted by Proteome Discoverer. MS/MS samples were analyzed using Sequest (XCorr Only) as described in Supplementary methods.

### PAR overlay assay

Peptides were ordered from Genscript (see sequences in Supplementary methods) and diluted to 0.5 mg/mL in PBS. One microgram of each peptide was slot-blotted onto nitrocellulose membrane. Membranes were fixed in 2.5% glutaraldehyde/PBS for 30 min then washed with TBS-T (50 mM Tris-HCl, pH 7.5, 150 mM NaCl, 0.1% Tween-20), followed by incubation with

0.2 μM PAR polymer (Trevigen) in TBS-T for 30 min, three washes with excess TBS-T, and blocking with 5% milk in TBS-T for 15 min. PAR was detected using MABE1016 (1:1000 in TBS- T + 5% milk) and anti-rabbit-HRP (1:50000 in TBS-T + 5% milk). Membranes were then incubated with 0.5% Ponceau in 3% acetic acid for 10 min and washed in dH_2_O.

### Protein purification

Huntingtin and HAP40 constructs used in this study have been described previously[96] and in the Supplementary methods, and are available through Addgene.

### Fluorescence polarization

FAM-labeled 11-mer and 26-mer PAR were produced as described [97]. Fluorescence polarization reactions were carried out in 20 mM HEPES pH 7.4, 50 mM KCl, 2.5% glycerol, 1 mM TCEP, 2 mM MgCl_2_, and 0.005% Tween-20 with a final concentration of 25 nM PAR substrate and 10 µM protein in a total reaction volume of 10 µL. Reactions were analysed using a Synergy H4 plate reader (Biotek) and data analysed using Graphpad Prism (version 9.4.1).

### *In vitro* PARP1 reaction and atomic force microscopy

For analysis by western blot, 10 fmol recombinant PARP1 was incubated in a 15-uL reaction (5 mM NAD+, 1X activated DNA, 50 mM Tris HCl pH 7.75, 50 mM NaCl) with the indicated amounts of recombinant HTT-HAP40 for 2 hours at 30°C. Reactions were separated by SDS-PAGE and immunoblotted with MABE1016 pan-ADP-ribose detection reagent. Signal intensities were quantified using ImageJ. For analysis by AFM, 35 nM PARP1 was incubated with 5 μg/mL sonicated salmon sperm DNA (Abcam ab229278) in deposition buffer (12.5 mM HEPES pH 8, 12.5 mM KCl, 1 mM DTT) in the the presence of 100 μM NAD^+^ and 10LmM MgCl_2_ with or without 11.5 nM HTT-HAP40 at 37°C for 30 min. Samples were diluted 10-fold in deposition buffer containing 3 mM MgCl_2_ and 10 μL deposited on freshly cleaved mica for 5 min before rinsing with HPLC-grade water and drying under nitrogen stream. AFM images were captured in air on a Bruker Dimension iCon (Bruker, Santa-Barbara, CA, USA) in soft tapping mode with ScanAsyst-Air tip (Bruker). In this experiment, continuous force–distance curves were recorded at 256 x 256 pixels at line rate 0.996 Hz, and the tip was oscillated in the vertical direction with an amplitude of 100–300 nm and at low frequency (1–2 kHz). Images were created using Nanoscope Analysis version 2.0 (Bruker Corporation).

### Statistical analysis

#### Human CSF analyses

Statistical analysis was performed using STATA MP Version 18. P values of <0.05 were considered significant. CSF PAR values were assessed for normality and subsequently log transformed. We assessed potential confounders including age, gender, blood contamination and time in the freezer. Hemoglobin was used as an indicator of blood contamination. There was an association between CSF PAR and storage time in the freezer (**Fig S12**), therefore we included storage time as a covariate in all analyses. Group comparisons were assessed using multiple regression adding storage time as a covariate followed by post-estimation Wald tests (also known as a Wald Chi-Square test, used to assess the significance of the coefficients in a regression model). Bonferroni correction was used to adjust for multiple comparisons. Correlations were assessed using Pearson’s partial correlation.

#### Experiment-based analyses

All data are represented as mean ± SEM with 3 independent experiments unless otherwise stated in the figure legends. Statistical analysis was performed using GraphPad Prism Version 9.4.1 and described for each experiment in figure legends. Differences among multiple means were assessed by ANOVA followed by Tukey’s post hoc test. Non-normally distributed data were analyzed with nonparametric test (Mann-Whitney test). Assessments with P < 0.05 were considered significant. Eta squared values were used to determine gene status effect size, with 0.01 representing a small effect, 0.06 representing a moderate effect, and 0.14 representing a large effect[98].

## Acknowledgements

The authors gratefully acknowledge the participation of HD patients and families, without whom this work could not be completed. This research was supported by the Canadian Institutes of Health Research Project Grant (MOP-119391) and the Krembil Foundation (RT), the Huntington Disease Society of America Berman Topper Career Development Fellowship and HD Human Biology Project (TM), Canadian Institutes of Health Research Project Grant (PJT180258 to SNA).

This work was also supported, in part, by a grant from the NIH R37 NS067525. T.M.D. is the Leonard and Madlyn Abramson Professor in Neurodegenerative Diseases. Support also included grants from the NIH to AKLL (R01GM104135), MD (T32GM080189), and MB (T32- CA009110), and from NINDS to LMT (R35 NS116872).

EJW reports research grants from Medical Research Council (MR/M008592/1), CHDI Foundation, European Huntington Disease Network, and F. Hoffmann-La Roche Ltd. The HD-CSF study was undertaken at the Leonard Wolfson Experimental Neurology Centre, University College London, supported by the National Institute for Health Research UCL Hospitals Biomedical Research Centre.

LMB is funded by the Medical research council (MR/W026686/1).

The Structural Genomics Consortium is a registered charity (no: 1097737) that receives funds from Bayer AG, Boehringer Ingelheim, Bristol Myers Squibb, Genentech, Genome Canada through Ontario Genomics Institute [OGI-196], EU/EFPIA/OICR/McGill/KTH/Diamond Innovative Medicines Initiative 2 Joint Undertaking [EUbOPEN grant 875510], Janssen, Merck KGaA (aka EMD in Canada and US), Pfizer and Takeda.

## Author contributions

TM conceived the project, designed and conducted experiments, analyzed and interpreted data, supervised the project and wrote the manuscript. CBB, RJH, LMB, FBR, and MD designed and conducted experiments, analyzed and interpreted data and contributed to drafting and editing the manuscript. NB, TK, MMW, KN, MM, MB, and KW conducted experiments and analyzed data. AKLL, SNA, EJW, TMD, VLD, CHA, LMT and RT supervised the work, analyzed and interpreted data and contributed to drafting and editing the manuscript.

## Materials and correspondence

All correspondence and requests for materials should be sent to TM or RT.

## References

1. Lee J-M, Ramos EM, Lee J-H, Gillis T, Mysore JS, Hayden MR, et al. CAG repeat expansion in Huntington disease determines age at onset in a fully dominant fashion. Neurology. 2012;78: 690–695.

2. Genetic Modifiers of Huntington’s Disease (GeM-HD) Consortium. CAG Repeat Not Polyglutamine Length Determines Timing of Huntington’s Disease Onset. Cell. 2019;178: 887–900.e14.

3. Genetic Modifiers of Huntington’s Disease (GeM-HD) Consortium. Identification of Genetic Factors that Modify Clinical Onset of Huntington’s Disease. Cell. 2015;162: 516–526.

4. Djousse L, Knowlton B, Hayden M, Almqvist EW, Brinkman R, Ross C, et al. Interaction of normal and expanded CAG repeat sizes influences age at onset of Huntington disease. Am J Med Genet A. 2003;119: 279–282.

5. Wexler NS, Lorimer J, Porter J, Gomez F, Moskowitz C, Shackell E, et al. Venezuelan kindreds reveal that genetic and environmental factors modulate Huntington’s disease age of onset. Proc Natl Acad Sci U S A. 2004;101: 3498–3503.

6. Bettencourt C, Hensman-Moss D, Flower M, Wiethoff S, Brice A, Goizet C, et al. DNA repair pathways underlie a common genetic mechanism modulating onset in polyglutamine diseases. Ann Neurol. 2016;79: 983–990.

7. Moss DJH, Pardiñas AF, Langbehn D, Lo K, Leavitt BR, Roos R, et al. Identification of genetic variants associated with Huntington’s disease progression: a genome-wide association study. Lancet Neurol. 2017;16: 701–711.

8. Ciosi M, Maxwell A, Cumming SA, Hensman Moss DJ, Alshammari AM, Flower MD, et al. A genetic association study of glutamine-encoding DNA sequence structures, somatic CAG expansion, and DNA repair gene variants, with Huntington disease clinical outcomes. EBioMedicine. 2019;48: 568–580.

9. Goold R, Flower M, Moss DH, Medway C, Wood-Kaczmar A, Andre R, et al. FAN1 modifies Huntington’s disease progression by stabilizing the expanded HTT CAG repeat. Hum Mol Genet. 2019;28: 650–661.

10. Flower M, Lomeikaite V, Ciosi M, Cumming S, Morales F, Lo K, et al. MSH3 modifies somatic instability and disease severity in Huntington’s and myotonic dystrophy type 1. Brain. 2019. doi:10.1093/brain/awz115

11. Lu X-H, Mattis VB, Wang N, Al-Ramahi I, van den Berg N, Fratantoni SA, et al. Targeting ATM ameliorates mutant Huntingtin toxicity in cell and animal models of Huntington’s disease. Sci Transl Med. 2014;6: 268ra178.

12. Maiuri T, Mocle AJ, Hung CL, Xia J, van Roon-Mom WMC, Truant R. Huntingtin is a scaffolding protein in the ATM oxidative DNA damage response complex. Hum Mol Genet. 2017;26: 395–406.

13 Askeland G, Dosoudilova Z, Rodinova M, Klempir J, Liskova I, Kuśnierczyk A, et al. Increased nuclear DNA damage precedes mitochondrial dysfunction in peripheral blood mononuclear cells from Huntington’s disease patients. Sci Rep. 2018;8: 9817.

14. Castaldo I, De Rosa M, Romano A, Zuchegna C, Squitieri F, Mechelli R, et al. DNA damage signatures in peripheral blood cells as biomarkers in prodromal huntington disease. Ann Neurol. 2019;85: 296–301.

15. Scudiero DA, Meyer SA, Clatterbuck BE, Tarone RE, Robbins JH. Hypersensitivity to N-methyl-N’-nitro-N-nitrosoguanidine in fibroblasts from patients with Huntington disease, familial dysautonomia, and other primary neuronal degenerations. Proc Natl Acad Sci U S A. 1981;78: 6451–6455.

16. Moshell AN, Tarone RE, Barrett SF, Robbins JH. Radiosensitivity in Huntington’s disease: implications for pathogenesis and presymptomatic diagnosis. Lancet. 1980;1: 9–11.

17. Ooi J, Langley SR, Xu X, Utami KH, Sim B, Huang Y, et al. Unbiased Profiling of Isogenic Huntington Disease hPSC-Derived CNS and Peripheral Cells Reveals Strong Cell-Type Specificity of CAG Length Effects. Cell Reports. 2019. pp. 2494–2508.e7. doi:10.1016/j.celrep.2019.02.008

18. Palminha NM, Dos Santos Souza C, Griffin J, Liao C, Ferraiuolo L, El-Khamisy SF. Defective repair of topoisomerase I induced chromosomal damage in Huntington’s disease. Cell Mol Life Sci. 2022;79: 1–21.

19. Lange J, Gillham O, Flower M, Ging H, Eaton S, Kapadia S, Neueder A, Duchen MR, Ferretti P, Tabrizi SJ. PolyQ length-dependent metabolic alterations and DNA damage drive human astrocyte dysfunction in Huntington’s disease. Prog Neurobiol. 2023; 102448.

20. Ferlazzo ML, Sonzogni L, Granzotto A, Bodgi L, Lartin O, Devic C, et al. Mutations of the Huntington’s disease protein impact on the ATM-dependent signaling and repair pathways of the radiation-induced DNA double-strand breaks: corrective effect of statins and bisphosphonates. Mol Neurobiol. 2014;49. doi:10.1007/s12035-013-8591-7

21. Gao R, Chakraborty A, Geater C, Pradhan S, Gordon KL, Snowden J, et al. Mutant huntingtin impairs PNKP and ATXN3, disrupting DNA repair and transcription. Elife. 2019;8. doi:10.7554/eLife.42988

22. Morozko EL, Smith-Geater C, Monteys AM, Pradhan S, Lim RG, Langfelder P, et al. PIAS1 modulates striatal transcription, DNA damage repair, and SUMOylation with relevance to Huntington’s disease. Proc Natl Acad Sci U S A. 2021;118. doi:10.1073/pnas.2021836118

23. Lin X, Kapoor A, Gu Y, Chow MJ, Peng J, Zhao K, et al. Contributions of DNA Damage to Alzheimer’s Disease. Int J Mol Sci. 2020;21. doi:10.3390/ijms21051666

24. Gonzalez-Hunt CP, Sanders LH. DNA damage and repair in Parkinson’s disease: Recent advances and new opportunities. J Neurosci Res. 2020. doi:10.1002/jnr.24592

25. Massey TH, Jones L. The central role of DNA damage and repair in CAG repeat diseases. Dis Model Mech. 2018;11. doi:10.1242/dmm.031930

26. Yau WY, O’Connor E, Sullivan R, Akijian L, Wood NW. DNA repair in trinucleotide repeat ataxias. FEBS J. 2018;285: 3669–3682.

27. Penndorf D, Witte OW, Kretz A. DNA plasticity and damage in amyotrophic lateral sclerosis. Neural Regeneration Res. 2018;13: 173–180.

28. He L, Liang J, Chen C, Chen J, Shen Y, Sun S, et al. C9orf72 functions in the nucleus to regulate DNA damage repair. Cell Death Differ. 2022;30: 716–730.

29. Moreira MC, Barbot C, Tachi N, Kozuka N, Uchida E, Gibson T, et al. The gene mutated in ataxia-ocular apraxia 1 encodes the new HIT/Zn-finger protein aprataxin. Nat Genet. 2001;29: 189–193.

30. Bras J, Alonso I, Barbot C, Costa MM, Darwent L, Orme T, et al. Mutations in PNKP cause recessive ataxia with oculomotor apraxia type 4. Am J Hum Genet. 2015;96: 474–479.

31. Takashima H, Boerkoel CF, John J, Saifi GM, Salih MAM, Armstrong D, et al. Mutation of TDP1 , encoding a topoisomerase I–dependent DNA damage repair enzyme, in spinocerebellar ataxia with axonal neuropathy. Nat Genet. 2002;32: 267–272.

32. Hoch NC, Hanzlikova H, Rulten SL, Tétreault M, Komulainen E, Ju L, et al. XRCC1 mutation is associated with PARP1 hyperactivation and cerebellar ataxia. Nature. 2017;541: 87–91.

33. Jaspers NG, Gatti RA, Baan C, Linssen PC, Bootsma D. Genetic complementation analysis of ataxia telangiectasia and Nijmegen breakage syndrome: a survey of 50 patients. Cytogenet Cell Genet. 1988;49: 259–263.

34. Maiuri T, Hung CLK, Suart C, Begeja N, Barba-Bazan C, Peng Y, et al. DNA Repair in Huntington’s Disease and Spinocerebellar Ataxias: Somatic Instability and Alternative Hypotheses. J Huntingtons Dis. 2021;10: 165–173.

35. Suart C, Perez AM, Al-Ramahi I, Maiuri T, Botas J, Truant R. Spinocerebellar Ataxia Type 1 protein Ataxin-1 is signalled to DNA damage by Ataxia Telangiectasia Mutated kinase. 2019. p. 701953. doi:10.1101/701953

36. Narne P, Pandey V, Simhadri PK, Phanithi PB. Poly(ADP-ribose)polymerase-1 hyperactivation in neurodegenerative diseases: The death knell tolls for neurons. Semin Cell Dev Biol. 2017;63: 154–166.

37. Shieh WM, Amé JC, Wilson MV, Wang ZQ, Koh DW, Jacobson MK, et al. Poly(ADP-ribose) polymerase null mouse cells synthesize ADP-ribose polymers. J Biol Chem. 1998;273: 30069–30072.

38. Amé JC, Rolli V, Schreiber V, Niedergang C, Apiou F, Decker P, et al. PARP-2, A novel mammalian DNA damage-dependent poly(ADP-ribose) polymerase. J Biol Chem. 1999;274: 17860–17868.

39. Masson M, Niedergang C, Schreiber V, Muller S, Menissier-de Murcia J, de Murcia G. XRCC1 is specifically associated with poly(ADP-ribose) polymerase and negatively regulates its activity following DNA damage. Mol Cell Biol. 1998;18: 3563–3571.

40. El-Khamisy SF, Masutani M, Suzuki H, Caldecott KW. A requirement for PARP-1 for the assembly or stability of XRCC1 nuclear foci at sites of oxidative DNA damage. Nucleic Acids Res. 2003;31: 5526–5533.

41. Kamaletdinova T, Fanaei-Kahrani Z, Wang Z-Q. The Enigmatic Function of PARP1: From PARylation Activity to PAR Readers. Cells. 2019;8. doi:10.3390/cells8121625

42. Wei L, Nakajima S, Hsieh C-L, Kanno S, Masutani M, Levine AS, et al. Damage response of XRCC1 at sites of DNA single strand breaks is regulated by phosphorylation and ubiquitylation after degradation of poly(ADP-ribose). J Cell Sci. 2013;126: 4414–4423.

43. Maruta H, Matsumura N, Tanuma S-I. Role of (ADP-ribose)nCatabolism in DNA Repair. Biochem Biophys Res Commun. 1997;236: 265–269.

44. Oei SL, Ziegler M. ATP for the DNA Ligation Step in Base Excision Repair Is Generated from Poly(ADP-ribose). J Biol Chem. 2000;275: 23234–23239.

45. Morales J, Li L, Fattah FJ, Dong Y, Bey EA, Patel M, et al. Review of poly (ADP-ribose) polymerase (PARP) mechanisms of action and rationale for targeting in cancer and other diseases. Crit Rev Eukaryot Gene Expr. 2014;24: 15–28.

46. Fatokun AA, Dawson VL, Dawson TM. Parthanatos: mitochondrial-linked mechanisms and therapeutic opportunities. Br J Pharmacol. 2014;171: 2000–2016.

47. Park H, Kam T-I, Dawson TM, Dawson VL. Poly (ADP-ribose) (PAR)-dependent cell death in neurodegenerative diseases. Int Rev Cell Mol Biol. 2020;353: 1–29.

48. Love S, Barber R, Wilcock GK. Increased poly(ADP-ribosyl)ation of nuclear proteins in Alzheimer’s disease. Brain. 1999. pp. 247–253. doi:10.1093/brain/122.2.247

49. McGurk L, Mojsilovic-Petrovic J, Van Deerlin VM, Shorter J, Kalb RG, Lee VM, et al. Nuclear poly(ADP-ribose) activity is a therapeutic target in amyotrophic lateral sclerosis. Acta Neuropathologica Communications. 2018;6: 84.

50. Kam T-I, Mao X, Park H, Chou S-C, Karuppagounder SS, Umanah GE, et al. Poly(ADP-ribose) drives pathologic α-synuclein neurodegeneration in Parkinson’s disease. Science. 2018;362. doi:10.1126/science.aat8407

51. Cardinale A, Paldino E, Giampà C, Bernardi G, Fusco FR. PARP-1 Inhibition Is Neuroprotective in the R6/2 Mouse Model of Huntington’s Disease. PLoS One. 2015;10: e0134482.

52. Paldino E, Cardinale A, D’Angelo V, Sauve I, Giampà C, Fusco FR. Selective Sparing of Striatal Interneurons after Poly (ADP-Ribose) Polymerase 1 Inhibition in the R6/2 Mouse Model of Huntington’s Disease. Front Neuroanat. 2017;11: 61.

53. Vis JC, Schipper E, de Boer-van Huizen RT, Verbeek MM, de Waal RMW, Wesseling P, et al. Expression pattern of apoptosis-related markers in Huntington’s disease. Acta Neuropathologica. 2005. pp. 321–328. doi:10.1007/s00401-004-0957-5

54. Thapa K, Khan H, Sharma U, Grewal AK, Singh TG. Poly (ADP-ribose) polymerase-1 as a promising drug target for neurodegenerative diseases. Life Sci. 2021;267: 118975.

55. Berger NA, Besson VC, Boulares AH, Bürkle A, Chiarugi A, Clark RS, et al. Opportunities for the repurposing of PARP inhibitors for the therapy of non-oncological diseases. Br J Pharmacol. 2018;175: 192–222.

56. Byrne LM, Rodrigues FB, Johnson EB, Wijeratne PA, De Vita E, Alexander DC, et al. Evaluation of mutant huntingtin and neurofilament proteins as potential markers in Huntington’s disease. Sci Transl Med. 2018;10. doi:10.1126/scitranslmed.aat7108

57. Rodrigues FB, Byrne LM, Tortelli R, Johnson EB, Wijeratne PA, Arridge M, et al. Mutant huntingtin and neurofilament light have distinct longitudinal dynamics in Huntington’s disease. Sci Transl Med. 2020;12. doi:10.1126/scitranslmed.abc2888

58. Hung CL-K, Maiuri T, Bowie LE, Gotesman R, Son S, Falcone M, et al. A Patient-Derived Cellular Model for Huntington’s Disease Reveals Phenotypes at Clinically Relevant CAG Lengths. Mol Biol Cell. 2018; mbcE18090590.

59. Holton NW, Ebenstein Y, Gassman NR. Broad spectrum detection of DNA damage by Repair Assisted Damage Detection (RADD). DNA Repair . 2018;66–67: 42–49.

60. James DI, Durant S, Eckersley K, Fairweather E, Griffiths LA, Hamilton N, et al. An assay to measure poly(ADP ribose) glycohydrolase (PARG) activity in cells. F1000Res. 2016;5: 736.

61. Kanev P-B, Varhoshkova S, Georgieva I, Lukarska M, Kirova D, Danovski G, et al. A unified mechanism for PARP inhibitor-induced PARP1 chromatin retention at DNA damage sites in living cells. Cell Rep. 2024;43: 114234.

62. Xue H, Bhardwaj A, Yin Y, Fijen C, Ephstein A, Zhang L, et al. A two-step mechanism governing PARP1-DNA retention by PARP inhibitors. Sci Adv. 2022;8: eabq0414.

63. Caldecott KW. XRCC1 protein; Form and function. DNA Repair . 2019;81. doi:10.1016/j.dnarep.2019.102664

64. Gagné J-P, Isabelle M, Lo KS, Bourassa S, Hendzel MJ, Dawson VL, et al. Proteome-wide identification of poly(ADP-ribose) binding proteins and poly(ADP-ribose)-associated protein complexes. Nucleic Acids Res. 2008;36: 6959–6976.

65. Zhang Y, Wang J, Ding M, Yu Y. Site-specific characterization of the Asp- and Glu-ADP-ribosylated proteome. Nat Methods. 2013;10: 981–984.

66. Jungmichel S, Rosenthal F, Altmeyer M, Lukas J, Hottiger MO, Nielsen ML. Proteome-wide identification of poly(ADP-Ribosyl)ation targets in different genotoxic stress responses. Mol Cell. 2013;52: 272–285.

67. Kanai Y. Overview on poly(ADP-ribose) immuno-biomedicine and future prospects. Proc Jpn Acad Ser B Phys Biol Sci. 2016;92: 222.

68. Kanai Y, Tanuma S, Sugimura T. Immunofluorescent staining of poly(ADP-ribose) in situ in HeLa cell chromosomes in the M phase. Proc Natl Acad Sci U S A. 1981;78: 2801.

69. Atwal RS, Desmond CR, Caron N, Maiuri T, Xia J, Sipione S, et al. Kinase inhibitors modulate huntingtin cell localization and toxicity. Nat Chem Biol. 2011;7: 453–460.

70. Godin JD, Colombo K, Molina-Calavita M, Keryer G, Zala D, Charrin BC, et al. Huntingtin is required for mitotic spindle orientation and mammalian neurogenesis. Neuron. 2010;67: 392–406.

71. Pleschke JM, Kleczkowska HE, Strohm M, Althaus FR. Poly (ADP-ribose) binds to specific domains in DNA damage checkpoint proteins. J Biol Chem. 2000;275: 40974–40980.

72. Gagné J-P, Hunter JM, Labrecque B, Chabot B, Poirier GG. A proteomic approach to the identification of heterogeneous nuclear ribonucleoproteins as a new family of poly(ADP-ribose)-binding proteins. Biochem J. 2003;371: 331–340.

73. Harding RJ, Deme JC, Hevler JF, Tamara S, Lemak A, Cantle JP, et al. Huntingtin structure is orchestrated by HAP40 and shows a polyglutamine expansion-specific interaction with exon 1. Communications Biology. 2021;4: 1–16.

74. Xiao J, Liu M, Qi Y, Chaban Y, Gao C, Pan B, et al. Structural insights into the activation of ATM kinase. Cell Res. 2019;29: 683–685.

75. Reber JM, Mangerich A. Why structure and chain length matter: on the biological significance underlying the structural heterogeneity of poly(ADP-ribose). Nucleic Acids Res. 2021;49: 8432–8448.

76 Alteen MG, Deme JC, Alvarez CP, Loppnau P, Hutchinson A, Seitova A, et al. Expanding the Huntingtons disease research toolbox; validated huntingtin subdomain constructs for biochemical and structural investigation of the huntingtin protein. bioRxiv. 2022. p. 2022.11.21.516512. doi:10.1101/2022.11.21.516512

77. Naumenko KN, Sukhanova MV, Hamon L, Kurgina TA, Anarbaev RO, Mangerich A, et al. The C-Terminal Domain of Y-Box Binding Protein 1 Exhibits Structure-Specific Binding to Poly(ADP-Ribose), Which Regulates PARP1 Activity. Frontiers in Cell and Developmental Biology. 2022;10. doi:10.3389/fcell.2022.831741

78. Sukhanova MV, Abrakhi S, Joshi V, Pastre D, Kutuzov MM, Anarbaev RO, et al. Single molecule detection of PARP1 and PARP2 interaction with DNA strand breaks and their poly(ADP-ribosyl)ation using high-resolution AFM imaging. Nucleic Acids Res. 2015;44: e60–e60.

79. Naumenko KN, Sukhanova MV, Hamon L, Kurgina TA, Alemasova EE, Kutuzov MM, et al. Regulation of Poly(ADP-Ribose) Polymerase 1 Activity by Y-Box-Binding Protein 1. Biomolecules. 2020;10: 1325.

80. Saudou F, Humbert S. The Biology of Huntingtin. Neuron. 2016;89: 910–926.

81. Desmond CR, Atwal RS, Xia J, Truant R. Identification of a karyopherin β1/β2 proline-tyrosine nuclear localization signal in huntingtin protein. J Biol Chem. 2012;287: 39626– 39633.

82. Xia J, Lee DH, Taylor J, Vandelft M, Truant R. Huntingtin contains a highly conserved nuclear export signal. Hum Mol Genet. 2003;12: 1393–1403.

83. Maiuri T, Woloshansky T, Xia J, Truant R. The huntingtin N17 domain is a multifunctional CRM1 and Ran-dependent nuclear and cilial export signal. Hum Mol Genet. 2013;22: 1383–1394.

84. Atwal RS, Xia J, Pinchev D, Taylor J, Epand RM, Truant R. Huntingtin has a membrane association signal that can modulate huntingtin aggregation, nuclear entry and toxicity. Hum Mol Genet. 2007;16: 2600–2615.

85. DiGiovanni LF, Mocle AJ, Xia J, Truant R. Huntingtin N17 domain is a reactive oxygen species sensor regulating huntingtin phosphorylation and localization. Hum Mol Genet. 2016. doi:10.1093/hmg/ddw234

86. Maiuri T, Bowie LE, Truant R. DNA Repair Signaling of Huntingtin: The Next Link Between Late-Onset Neurodegenerative Disease and Oxidative DNA Damage. DNA Cell Biol. 2019;38: 1–6.

87. Slade D. PARP and PARG inhibitors in cancer treatment. Genes Dev. 2020;34: 360–394.

88. Byrne LM, Wild EJ. Cerebrospinal Fluid Biomarkers for Huntington’s Disease. J Huntingtons Dis. 2016;5: 1–13.

89. McNulty P, Pilcher R, Ramesh R, Necuiniate R, Hughes A, Farewell D, et al. Reduced Cancer Incidence in Huntington’s Disease: Analysis in the Registry Study. J Huntingtons Dis. 2018;7: 209–222.

90. Rubinsztein DC, Amos W, Leggo J, Goodburn S, Ramesar RS, Old J, et al. Mutational bias provides a model for the evolution of Huntington’s disease and predicts a general increase in disease prevalence. Nat Genet. 1994;7: 525–530.

91. Landwehrmeyer GB, Fitzer-Attas CJ, Giuliano JD, Gonçalves N, Anderson KE, Cardoso F, et al. Data Analytics from Enroll-HD, a Global Clinical Research Platform for Huntington’s Disease. Mov Disord Clin Pract. 2017;4: 212–224.

92. Trettel F, Rigamonti D, Hilditch-Maguire P, Wheeler VC, Sharp AH, Persichetti F, et al. Dominant phenotypes produced by the HD mutation in STHdh(Q111) striatal cells. Hum Mol Genet. 2000;9: 2799–2809.

93. Smith-Geater C, Hernandez SJ, Lim RG, Adam M, Wu J, Stocksdale JT, et al. Aberrant Development Corrected in Adult-Onset Huntington’s Disease iPSC-Derived Neuronal Cultures via WNT Signaling Modulation. Stem Cell Reports. 2020;14: 406–419.

94. Goss S, Bazan CB, Neuman K, Peng C, Begeja N, Suart CE, et al. Mod3D: A low-cost, flexible modular system of live-cell microscopy chambers and holders. PLoS One. 2022;17: e0269345.

95. Carpenter AE, Jones TR, Lamprecht MR, Clarke C, Kang IH, Friman O, et al. CellProfiler: image analysis software for identifying and quantifying cell phenotypes. Genome Biol. 2006;7: R100.

96. Harding RJ, Loppnau P, Ackloo S, Lemak A, Hutchinson A, Hunt B, et al. Design and characterization of mutant and wildtype huntingtin proteins produced from a toolkit of scalable eukaryotic expression systems. J Biol Chem. 2019;294: 6986–7001.

97. Ando Y, Elkayam E, McPherson RL, Dasovich M, Cheng S-J, Voorneveld J, et al. ELTA: Enzymatic Labeling of Terminal ADP-Ribose. Mol Cell. 2019;73: 845–856.e5.

98. Pallant J. SPSS survival manual: A step by step guide to data analysis using IBM SPSS. [cited 11 Oct 2022]. doi:10.4324/9781003117452/spss-survival-manual-julie-pallant

