## Supplemental figures and methods for "Poly ADP-Ribose Signaling is Dysregulated in Huntington Disease"

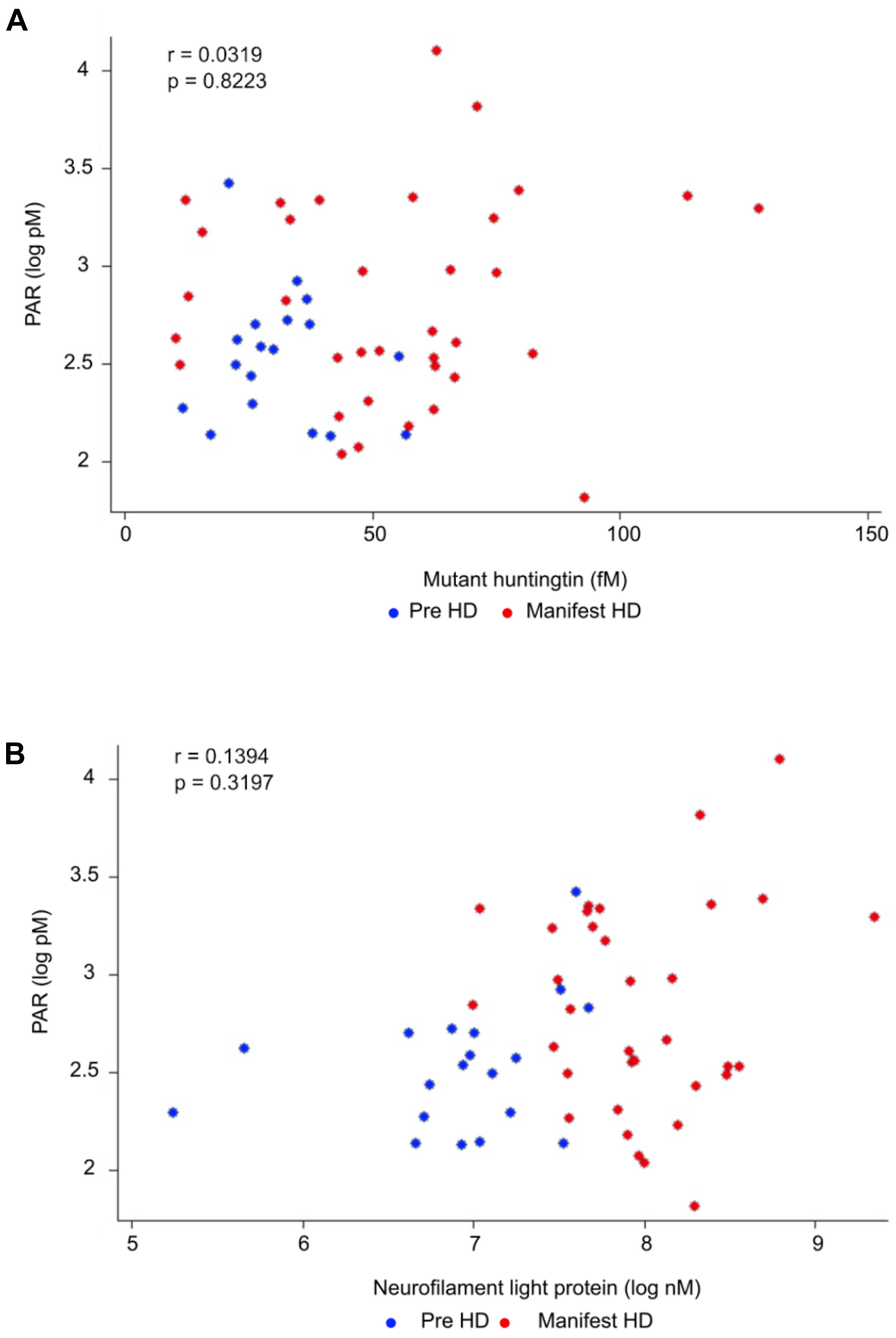


**Figure S1: CSF PAR levels do not correlate with HD biomarkers**
No association between CSF PAR and (A) mutant huntingtin or
(B) neurofilament light protein levels, assessed using partial correlations with time in freezer. CSF PAR is natural log transformed. CSF samples are from HD mutation carriers from the HD-CSF study (premanifest HD [n=35], and manifest HD [n=19]).

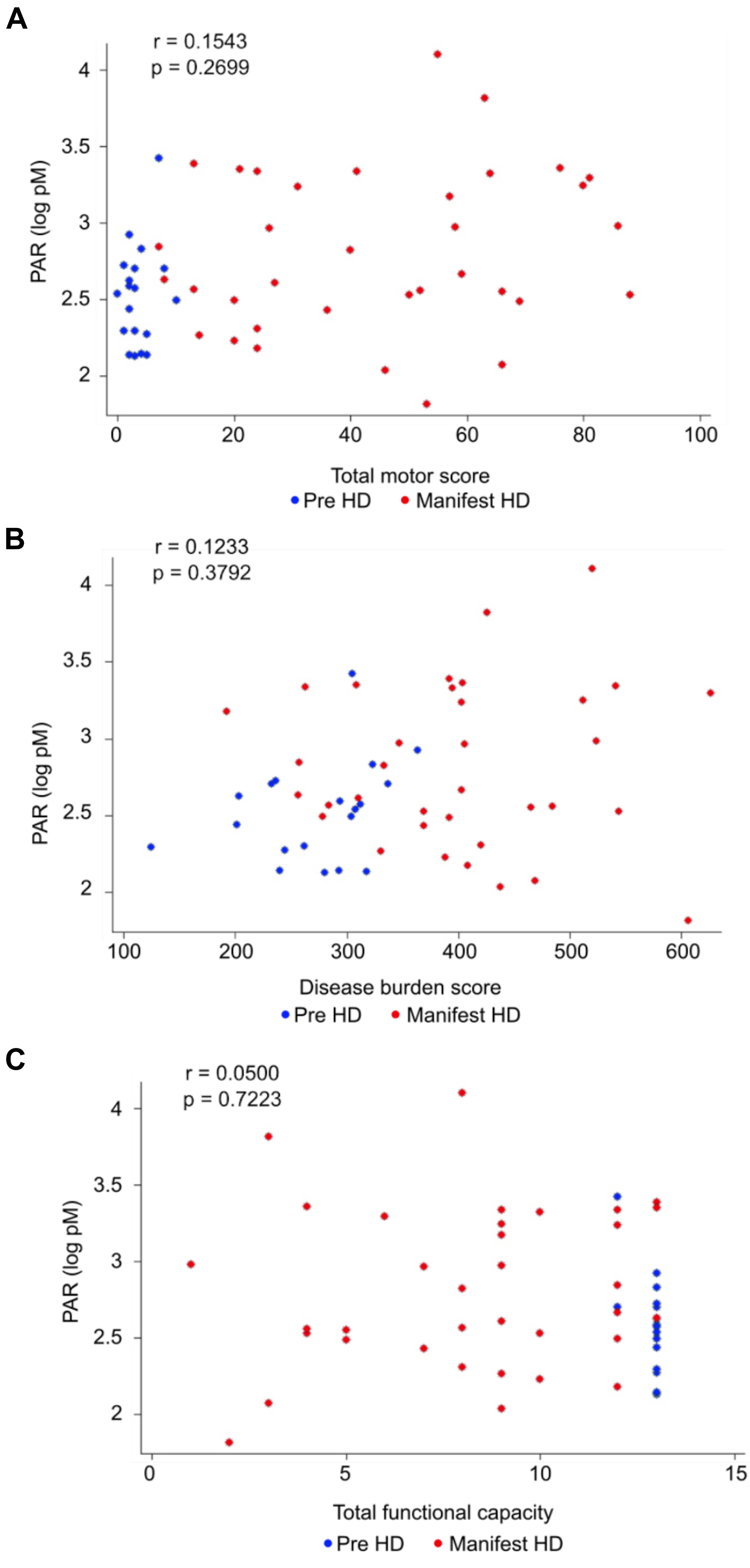


**Figure S2: CSF PAR levels do not correlate with clinical measures of disease progression**
No association between CSF PAR and UHDRS clinical scores including (A) total motor score, (B) disease burden score, and (C) total functional capacity, assessed using partial correlations with time in freezer. CSF PAR is natural log transformed. CSF samples are from HD mutation carriers from the HD-CSF study (premanifest HD [n=35], and manifest HD [n=19]).

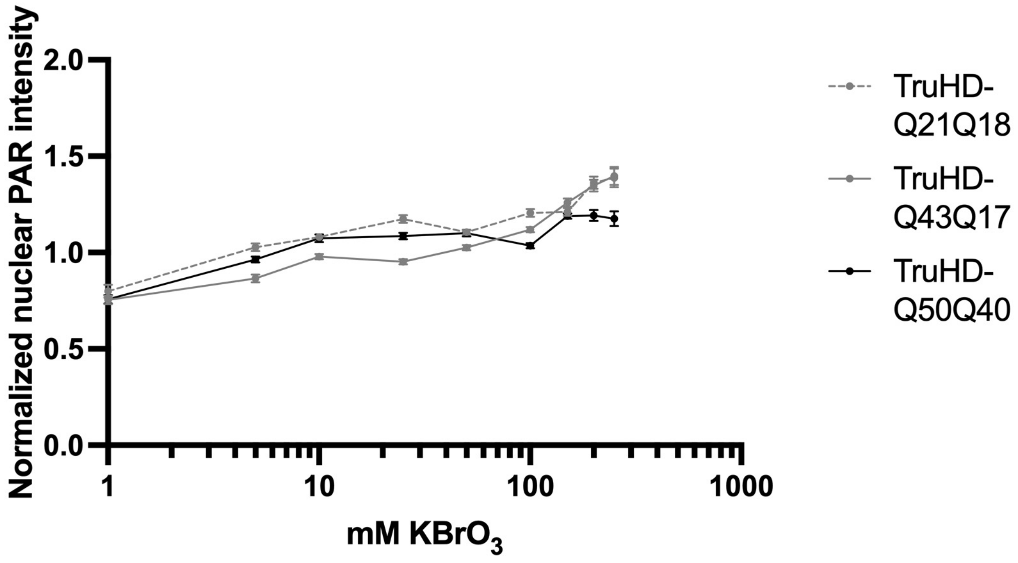


**Figure S3: PAR response in HD patient-derived fibroblasts**
TruHD cells were treated with increasing doses of $KBrO_3$ for 30 min and stained with MABE1031 PAR detection reagent. Nuclear PAR intensity was measured using CellProfiler, mean intensity recorded for each image (18 images per condition; >500 cells), and values normalized to the control condition. Data from three independent experiments is shown. Error bars: SEM.

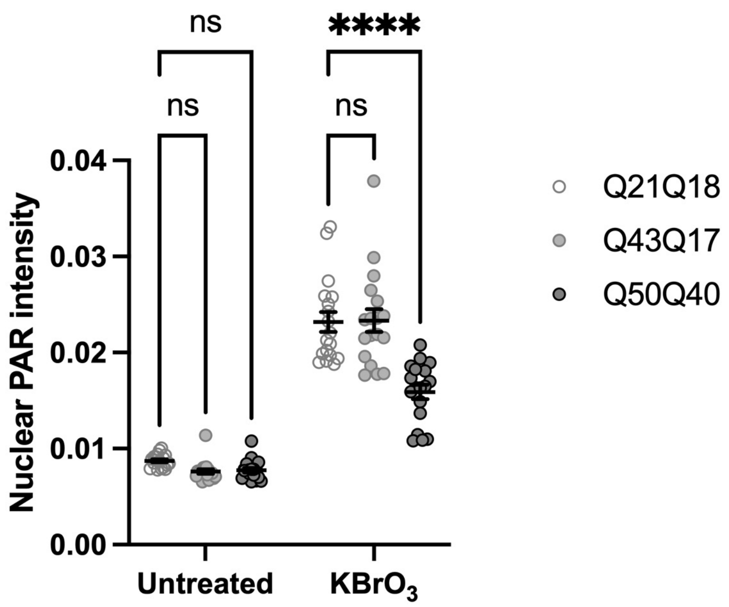


**Figure S4: Primary HD patient-derived fibroblasts are consistent with hTERT-immortalized cells**
Primary fibroblasts were treated with 100 mM $KBrO_3$ for 30 min followed by fixation and staining with MABE1031 PAR detection reagent. Nuclear PAR intensity was measured using CellProfiler, mean intensity recorded for each image (6 images per condition; >500 cells). Data from three experiments is shown. Results were analyzed by two-way ANOVA and corrected for multiple comparisons using Tukey's test. PAR intensity values for $KBrO_3$-treated cells were significantly higher than untreated cells ($p<0.0001$ for each cell line).

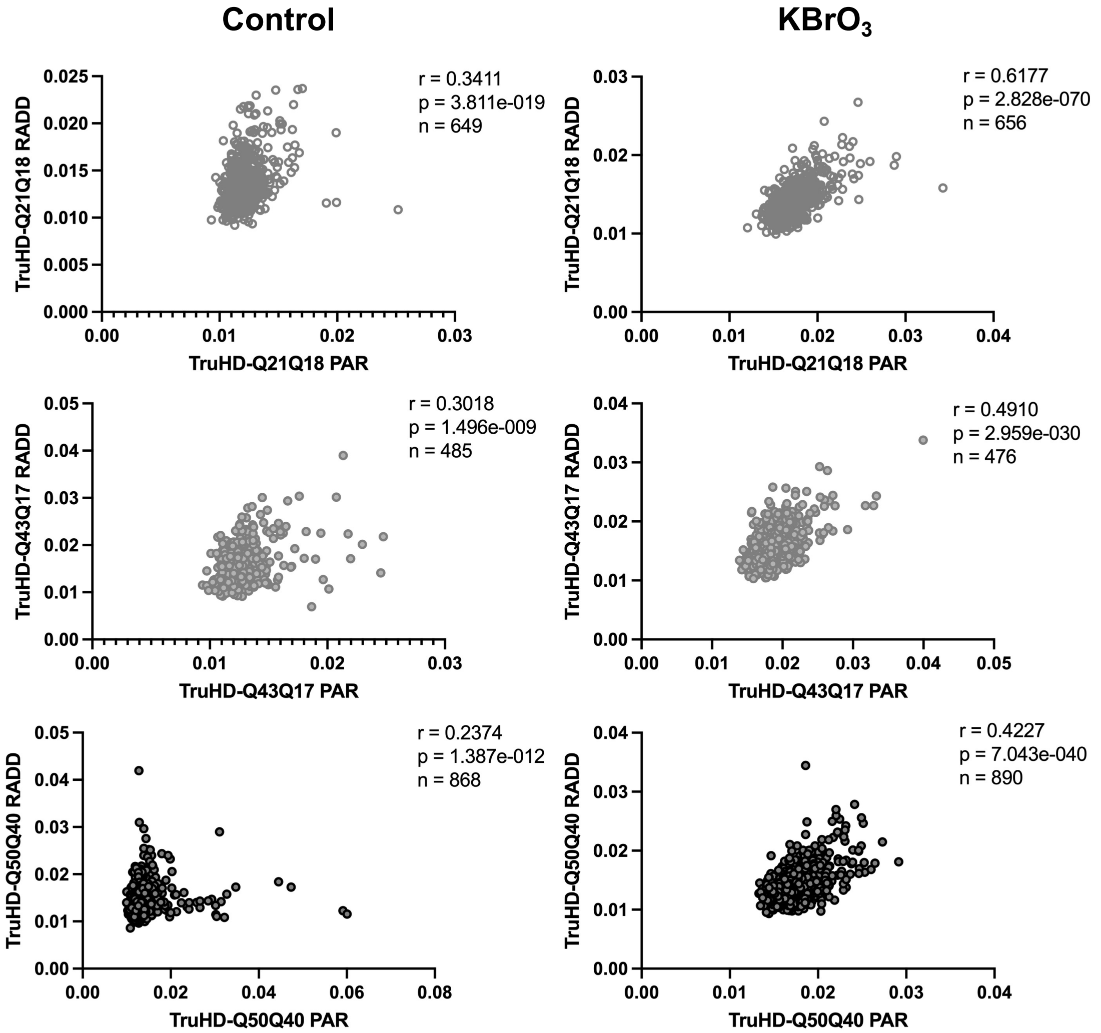


Trial 1

| Cell line | Spearman r value | |
|---|---|---|
| | Control | 100 mM KBrO3 |
| TruHD-Q21Q18 | 0.3411 | 0.6177 |
| TruHD-Q43Q17 | 0.3018 | 0.4910 |
| TruHD-Q50Q40 | 0.2374 | 0.4227 |

Trial 2

| Cell line | Spearman r value | |
|---|---|---|
| | Control | 100 mM KBrO3 |
| TruHD-Q21Q18 | 0.6554 | 0.7409 |
| TruHD-Q43Q17 | 0.4722 | 0.6031 |
| TruHD-Q50Q40 | 0.5438 | 0.7017 |

Trial 3

| Cell line | Spearman r value | |
|---|---|---|
| | Control | 100 mM KBrO3 |
| TruHD-Q21Q18 | 0.7015 | 0.8154 |
| TruHD-Q43Q17 | 0.5538 | 0.5692 |
| TruHD-Q50Q40 | 0.7274 | 0.6372 |

**Figure S5: Per-nucleus RADD:PAR correlation**

TruHD cells were treated with 100 mM KBrO3 for 30 min followed by Repair-Assisted Damage Detection (RADD) and co-staining with MABE1031 PAR detection reagent. Staining intensity within nuclei was quantified with CellProfiler and Spearman correlation coefficients calculated using GraphPad Prism. XY plots from trial 1 are shown. Table summarizes Spearman r values for three independent experiments.

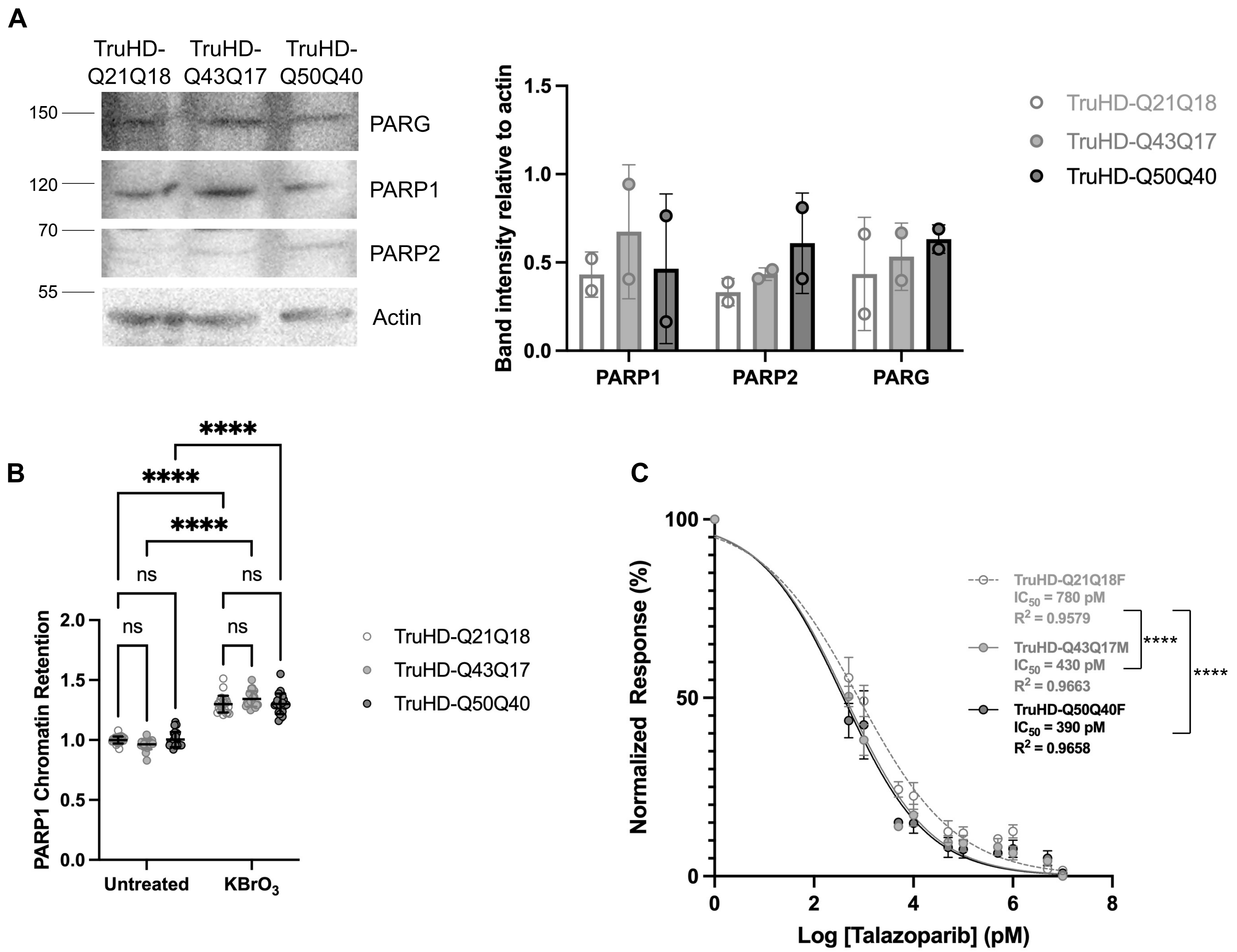


**Figure S6: PARP1/2 and PARG expression levels are similar in control and HD patient-derived cells**

**A**: Total protein lysates were separated by SDS-PAGE and probed with the indicated antibodies. Band intensities were quantified using ImageJ. Representative westerns and mean values from two independent experiments are shown. **B**: TruHD cells were treated with 100 mM $KBrO_3$ for 30 min and soluble proteins extracted with cold 0.2% Triton X-100 in PBS for 2 min on ice prior to fixation and permeabilization with cold methanol for 10 min at -20°C. The PARP1 retained on chromatin was visualized by immunofluorescence and quantified using CellProfiler. Data from two experiments is shown. Results were analyzed by two-way ANOVA and corrected for multiple comparisons using Tukey's test. PARP1 intensity values for $KBrO_3$-treated cells were significantly higher than untreated cells ($p<0.0001$ for each cell line). **C**: TruHD cells were treated with talazoparib as described in Fig 3B. Error bars = SEM for 5 experiments. **** $p<0.0001$ (Brown-Forsythe and Welch ANOVA tests).

| Database | Percentage of proteins found in other PARylated protein databases |
|---|---|
| PARylated proteins bound by 10H antibody | **11.8%** (57/483) |
| E- and D-PARylated proteins bound by boronate resin | **27.7%** (92/332) |
| PARylated proteins bound by macrodomain | **37.4%** (101/270) |

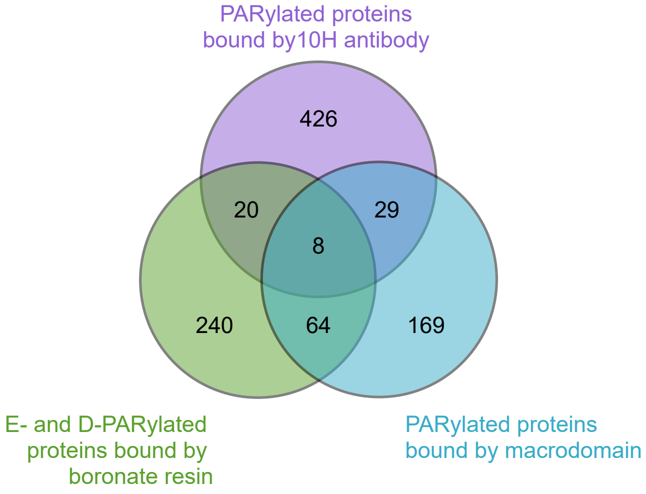


| Database | Percentage of proteins found in compiled list of PARylated proteins |
|---|---|
| Huntingtin interactors | **32%** (126/394) |

**Figure S7: Degree of overlap between databases of PARylated proteins**
Proteins from three independently produced databases of PARylated proteins were considered. The first database, generated by immunoprecipitation with the PAR-specific 10H antibody following alkylation-induced DNA damage of SK-N-SH (human neuroblastoma) cells, lists 482 PARylated proteins (Gagne et al, 2008). The second database was generated by alkylation-induced DNA damage of HCT116 (human colon carcinoma) cells followed by affinity chromatography with boronic acid-agarose, which forms an ester bond with a 1,2-cis-diol moiety of ADP-ribose. Elution by NH2OH leaves a hydroxamic acid derivative on aspartic acid and glutamic acid residues that can be distinguished by mass spectrometry. The resulting database lists 332 proteins representing the aspartic acid- and glutamic acid-ADP-ribosylated proteome (Zhang et al, 2013). The third database lists 270 proteins purified by the PAR-binding macrodomain, GST-Af1521, from U2OS (human bone osteosarcoma) cells after a panel of genotoxic stresses (Jungmichel et al, 2013). The different purification strategies yielded different sets of proteins, with some degree of overlap (right panel). The three databases were compiled and replicates removed to generate a final list of 955 PARylated proteins, which was compared to the list of huntingtin-interacting proteins (see Table S2).

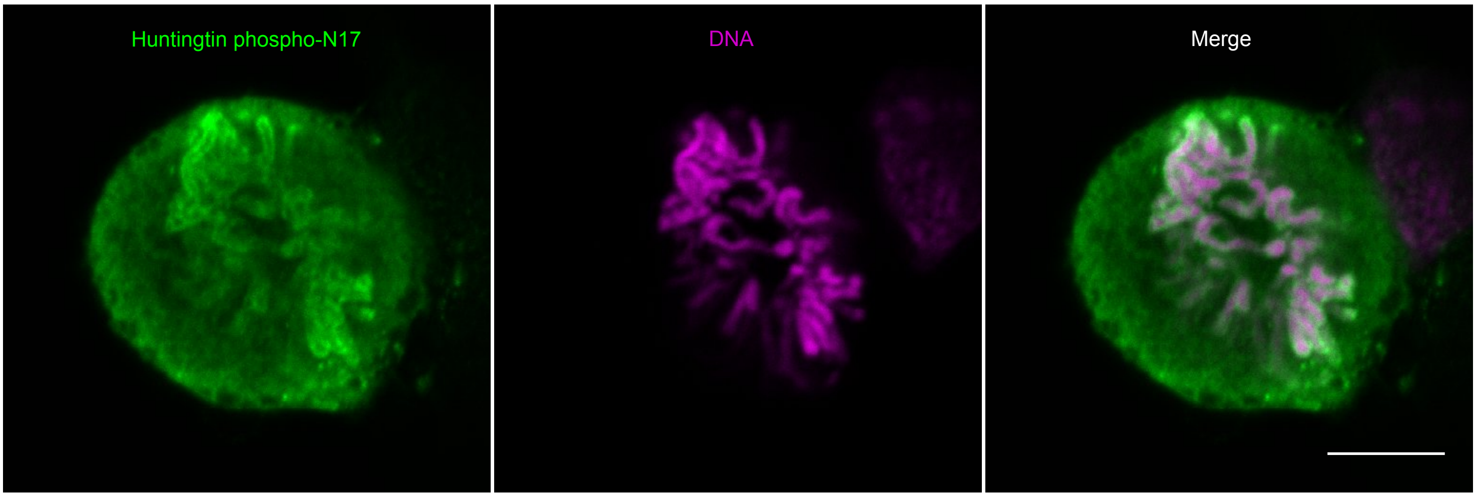


**Figure S8: Huntingtin localizes to mitotic chromosomes upon PAR accumulation**
RPE1 cells were treated with 10 µM PDD00017273 PARG inhibitor for 40 min prior to methanol fixation and immunofluorescence against huntingtin phosphorylated at residues S13 and S16 within the N17 domain. DNA was stained with Hoechst. Images are representative of all mitotic cells observed (n > 10 cells from two independent experiments).

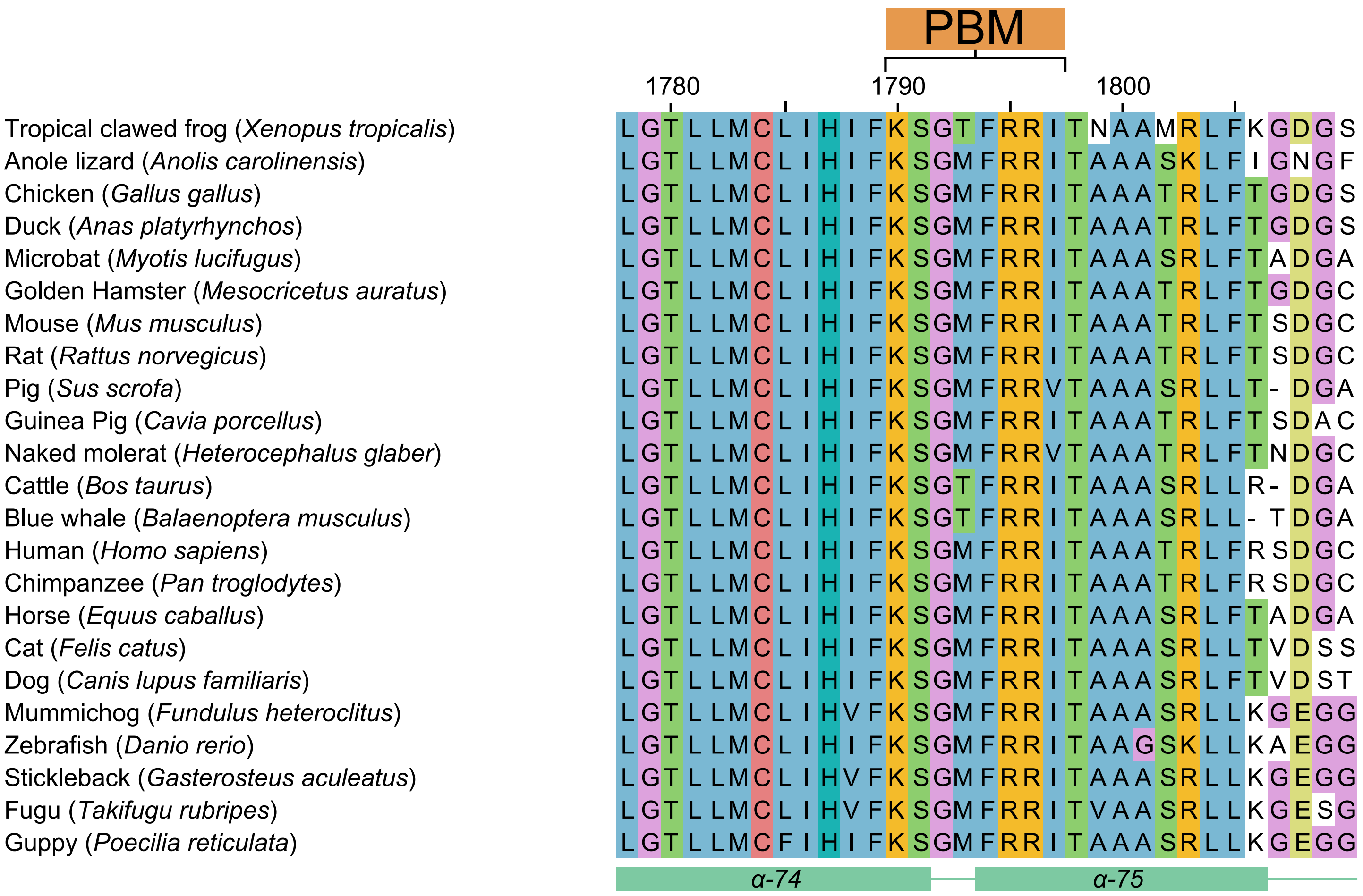


**Figure S9: The huntingtin PAR-binding motif is evolutionarily conserved**
Multiple sequence alignment generated by Clustal Omega and visualized with JalView of huntingtin from human and orthologous species extracted from Ensembl. PAR binding motif (PBM) and flanking regions are shown with numbering reflecting human Q23 sequence and secondary structure based on 6X9O high resolution cryoEM model.

A

B

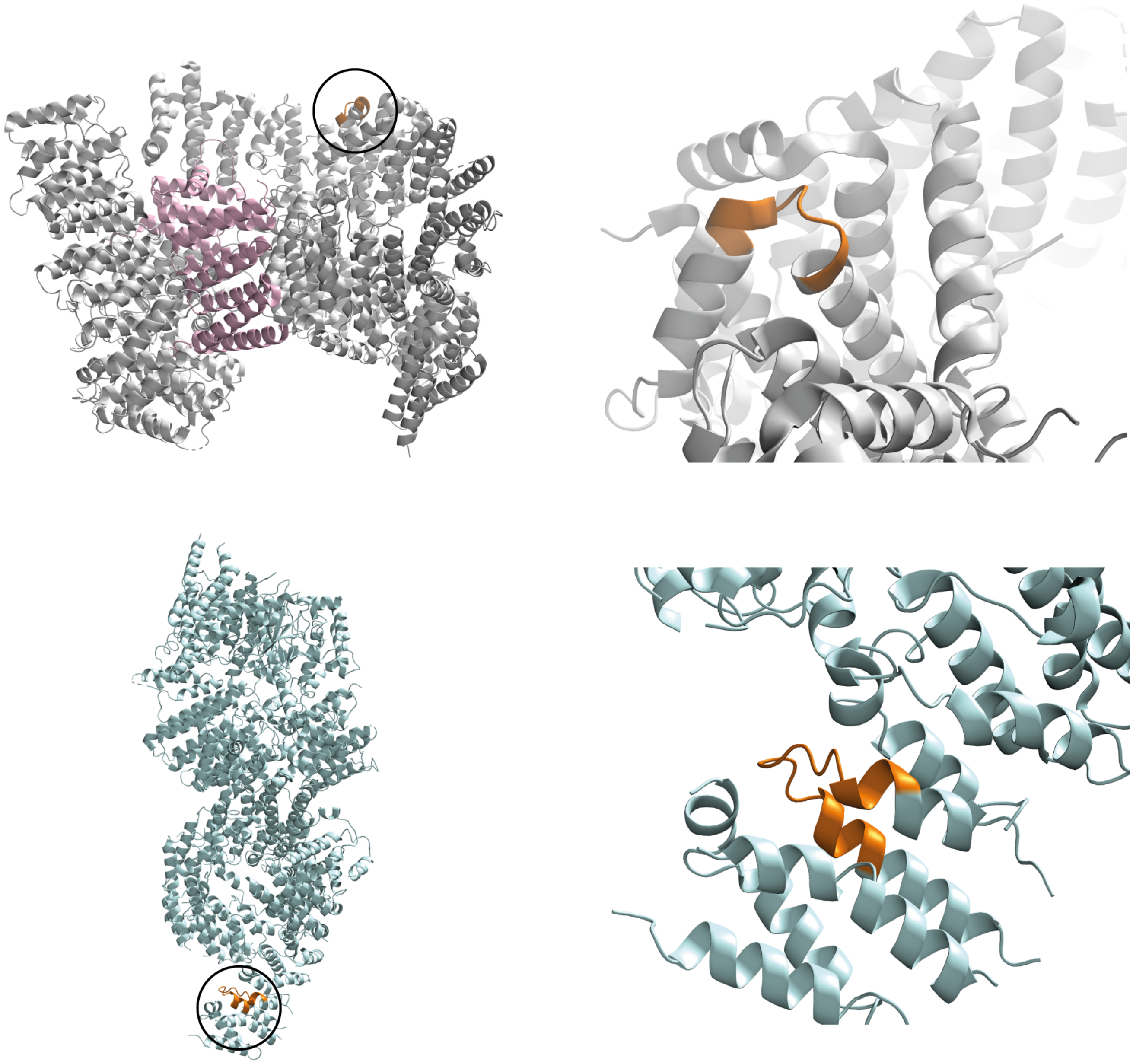

**Figure S10: The huntingtin PAR-binding motif is analogous to the Ataxia Telangiectasia Mutated (ATM) kinase PAR-binding site**

**A**: high resolution cryoEM model of huntingtin-HAP40 complex (PDB - 6X9O) shown in cartoon representation with huntingtin in grey, HAP40 in pink and the PAR-binding motif in orange (circled). The PAR-binding motif is a helix-turn-helix in a HEAT repeat in the bridge domain. **B**: mid-resolution cryoEM model of ATM kinase (PDB - 6K9L) shown in cartoon representation with ATM in cyan and the PAR-binding motif, located at the N-terminus of ATM, in orange (circled). Similar to huntingtin, the PAR-binding motif is a surface-exposed helix-turn-helix in a HEAT repeat.

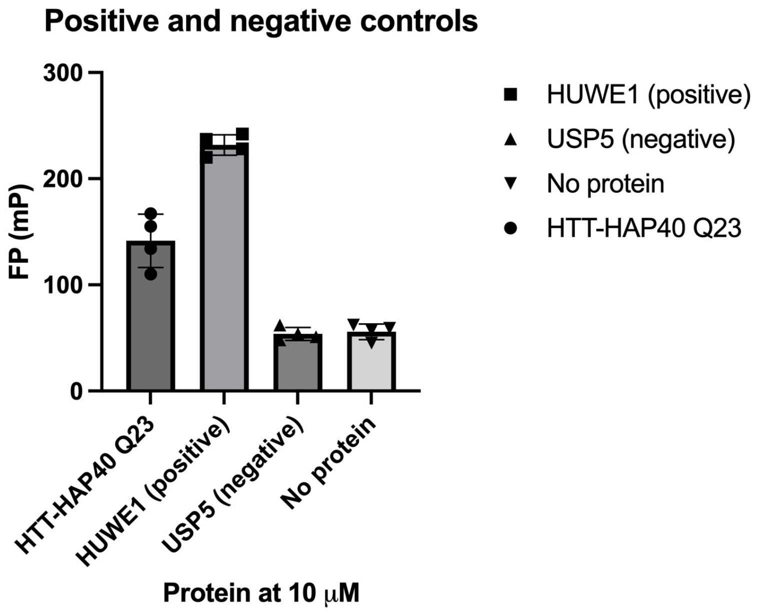


**Figure S11: Fluorescence polymerization positive and negative controls**
Fluorescence polarization assays using FAM-labeled 26-mer PAR and 10 μM of the indicated proteins. Reactions were carried out with 4 intra-assay replicates.

**A**

| | Healthy Controls | | | Premanifest HD | | | Manifest HD | | |
|---|---|---|---|---|---|---|---|---|---|
| | N | Mean | SD | N | Mean | SD | N | Mean | SD |
| **Age (yrs)** | 17 | 53.15 | 10.6 | 19 | 44.1 | 11.29 | 35 | 57.54 | 9.65 |
| **CAG** | NA | NA | NA | 19 | 41.89 | 1.595 | 35 | 42.69 | 2.311 |
| **DBS** | NA | NA | NA | 19 | 272.9 | 57.55 | 35 | 402 | 102.5 |
| **PAR (pM)** | 17 | 24.54 | 9.968 | 19 | 13.24 | 5.21 | 35 | 19.23 | 11.32 |
| **Storage time (days)** | 17 | 367.8 | 121.4 | 19 | 375.9 | 97.71 | 35 | 283.1 | 86.07 |
| **Hemoglobin (ng/mL)** | 15 | 535.7 | 488.1 | 17 | 272.6 | 230 | 31 | 432.7 | 376.2 |

SD, Standard Deviation; CAG, Cytosine-Adenine-Guanine repeat length number; NA, not applicable; DBS, Disease burden score; PAR, Poly ADP-ribose;

**B**

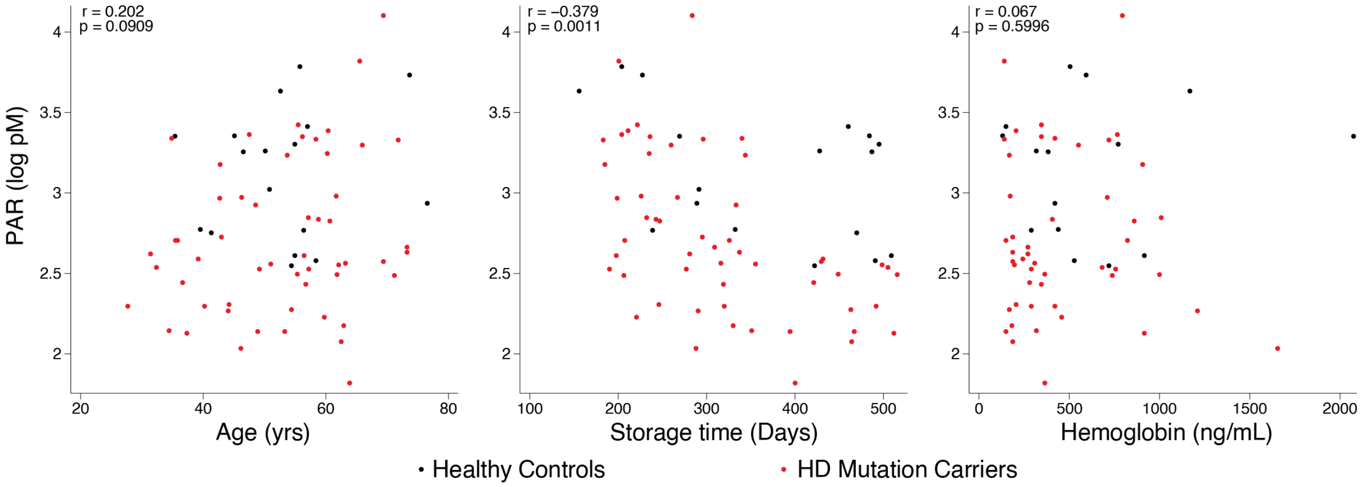


**Figure S12: Human CSF sample characteristics**

A: Participant characteristics for human CSF samples.

B: Assessment of potential confounding variables age, storage time in freezer, and blood contamination as measured by Hemoglobin. Healthy controls are in navy and HD mutation carriers in red. Statistics were generated by Pearson's correlation.

**Supplementary video 1: Huntingtin localizes to mitotic chromosomes upon PAR accumulation**
RPE1 cells were treated with 10 μM PDD00017273 PARG inhibitor for 40 min prior to methanol fixation and immunofluorescence against huntingtin phosphorylated at residues S13 and S16 within the N17 domain. DNA was stained with Hoechst. Z-stack images acquired on the Nikon A1+ confocal system were reconstructed for 3D video with the Nikon NIS-Elements Advanced Research 64-bit acquisition software. Video is representative of all mitotic cells observed (n > 10 cells from two independent experiments).

### Supplementary methods

**Antibodies**
PAR (MABE1031) and pan ADP-ribose (MABE1016) detection reagents were from MilliporeSigma. Rabbit monoclonal anti-huntingtin antibody (EPR5526) was from Abcam. Mouse monoclonal anti-PARP1 (611039) was from BD Biosciences. Rabbit polyclonal anti-PARP2 (AB_2793328) was from Active Motif. Mouse polyclonal anti-PARG (ab169639) was from Abcam. Rabbit polyclonal anti-actin (20-33) was from MilliporeSigma. Rabbit monoclonal anti-vinculin (ab129002) was from Abcam. Rabbit monoclonal anti-ɣH2AX (ab81299) was from Abcam. Rabbit polyclonal antibodies against huntingtin N17 domain were described and validated previously (Atwal et al, 2011). Anti-rabbit IgG and anti-mouse IgG antibodies conjugated to HRP were from Abcam. Anti-rabbit IgG and anti-mouse IgG antibodies conjugated to Alexa Fluor 488 or 594 were from Invitrogen.

**Repair assisted damage detection (RADD)**
Cells were incubated with cytoskeletal buffer (CSK, 100 mM NaCl, 300 mM sucrose, 10 mM PIPES pH 6.8, 3 mM $MgCl_2$, 0.5% Triton X-100) on ice for 5 min, followed by PBS wash and fixation in cold methanol for 10 min on ice. After PBS wash, cells were further permeabilized with 0.25% Triton X-100 in PBS for 10 min at room temperature, then washed with PBS followed by a wash with sterile deionized $H_2O$. The DNA damage processing mix contained Fapy-DNA glycosylase (NEB M0240S), Endo IV (NEB M0304S), and Endo VIII (NEB M0299S) prepared in 1x Thermpol buffer (NEB B9004S) containing 500 μM NAD+ and 200 μg/mL BSA and incubated at 37°C in a humidified incubator for one hour. The gap filling mix containing large Klenow fragment DNA polymerase I (NEB M0210S) and biotin-11-dUTP (ThermoFisher R0081) was added directly to the lesion processing mix and incubated for an additional hour. After washing with PBS, non-specific binding was blocked with 10% FBS in PBS for 30 min at room temperature followed by incubation with MABE1031 (1:1000 in blocking buffer) for one hour at room temperature. Chicken anti-rabbit Alexa Fluor 594 secondary antibody 1:500 in blocking buffer) was co-incubated with mouse monoclonal Alexa Fluor 488 conjugated anti-biotin antibody (BK-1/39, ThermoFisher 53-9895-82) (1:1000 in blocking buffer) for 30 min at room temperature, and cells were counterstained with Hoechst.

**Measurement of PAR levels in cells**
After the indicated treatments, cells were fixed and permeabilized with cold methanol for 10 min at -20°C. Fixed cells were washed with PBS, blocked with 10% FBS in PBS for 15 min and stained with either

MABE1031 (PAR-binding) or MABE1016 (pan-ADP-ribose-binding) reagent diluted 1:500 in blocking buffer for 1 hour at room temperature. Following two PBS washes, cells were incubated with anti-rabbit secondary antibody diluted 1:500 in blocking buffer for 15 min at room temperature, washed with PBS and incubated with 0.2 ug/mL Hoechst in PBS for 5 min. Cells were imaged in PBS using the Nikon A1+ confocal system on a Nikon Eclipse Ti inverted microscope equipped with GaAsP detectors, using a PLAN APO 20×/0.75 dry objective. Nuclei were identified as primary objects in CellProfiler using the Hoechst staining, then pixel intensity of the PAR staining within nuclei was calculated and the mean intensity recorded for each image. Ten to twelve images per well were captured, representing 800-1000 cells per experiment.

**Inhibitor dose response experiments**

TruHD cells were grown in CellCarrier-96 Ultra Microplates (Perkin Elmer) and pre-treated with a dilution series of veliparib or PDD00017273 (8 points from 0-10 μM in duplicate) in fresh growth media for one hour. Cells were then treated for 30 min with 100 mM $KBrO_3$ dissolved in PBS containing $Ca^{2+}$ and $Mg^{2+}$ in the presence of the 8-point inhibitor dose range. Veliparib or talazoparib pre-treatment and $KBrO_3$ treatments included 5 μM PARG inhibitor for all conditions to enable visualization of the pan-ADP-ribose signal by MABE1016. After treatment, cells were fixed in cold methanol at -20°C for 10 min, washed with PBS, and blocked in 10% FBS in PBS for 15 min at room temperature. Indirect immunofluorescence against pan-ADP-ribose was performed using MABE1016 (1:750 dilution in blocking buffer) for 30 min at room temperature followed by Alexa Fluor donkey anti-rabbit-488 (1:1000 dilution in blocking buffer) for 15 min at room temperature. Nuclei were counterstained with 0.2 ug/mL Hoechst for 5 min and cells were imaged in PBS. Six fields per well were imaged using the 10× objective on the EVOS FL Auto 2 widefield microscope (1000-2000 cells per condition). Using CellProfiler, nuclei were identified by the Hoechst signal, pan-ADP-ribose signal intensity within nuclei was calculated, and the mean nuclear signal intensities for each field were averaged for each condition.

**Immunofluorescence**

PARP1 chromatin retention assay: after extraction of soluble proteins as described in the main text, cells were fixed and permeabilized with cold methanol for 10 min at -20°C. Fixed cells were washed with PBS, blocked with 10% FBS in PBS for 15 min and stained with mouse anti-PARP1 antibody (BD Biosciences) diluted 1:500 in blocking buffer for 1 hour at room temperature. Following two PBS washes, cells were incubated with anti-mouse secondary antibody diluted 1:500 in blocking buffer for 15 min at room temperature, washed with PBS and incubated with 0.2 ug/mL Hoechst in PBS for 5 min. Cells were imaged in PBS using the 10× objective on the EVOS FL Auto 2 widefield microscope (12 fields of view per condition; 2000-3000 cells).

For imaging of mitotic cells, RPE1 cells were fixed in cold methanol for 20 min at -20°C and incubated with blocking buffer (10% FBS in PBS) for 15 min at room temperature followed by anti-huntingtin phospho S13S16 antibody (Atwal et al, 2011) (1:500 dilution in blocking buffer) for 45 min at room temperature, two washes with PBS, anti-rabbit secondary antibody (Alexa 594 or Alexa 488; 1:500 dilution in blocking buffer) for 20 min at room temperature and two washes with PBS. For co-staining of PARP1, cells were incubated with mouse anti-PARP1 antibody (BD Biosciences, 1:500 dilution in blocking buffer) for 1 hour at room temperature followed by two washes with PBS, anti-mouse secondary antibody (Alexa 488; 1:500 dilution in blocking buffer) for 20 min at room temperature and two washes with PBS. DNA was stained with 0.2 ug/mL Hoechst for 5 min at room temperature followed by a final PBS wash. Cell were imaged in PBS on a Nikon A1+ confocal system on a Nikon Eclipse Ti inverted microscope equipped with GaAsP detectors, using a PLAN APO 60×/1.40 oil objective. Image acquisition and Z-stack reconstruction was done with the Nikon NIS-Elements Advanced Research 64-bit acquisition software (Nikon).

For staining of iPSC-derived neurons, cells were grown on glass coverslips, fixed with 4% PFA, and stained as described above (Measurement of PAR levels in cells). Cells were co-stained with neuronal marker Map2 (Millipore MAB3418, 1:1000) and anti-mouse secondary antibody, and imaged using the 20× objective on the EVOS FL Auto 2 widefield microscope. Using CellProfiler, neurons were identified using the Map2 signal, nuclei were identified by the Hoechst signal, and PAR signal intensity within nuclei of neurons was measured.

**Cell lysis and western analysis**
Proteins were separated by SDS-PAGE using precast 4-20% gradient gels (Bio-Rad) and transferred to PVDF membrane (Millipore). Membranes were blocked in TBS-T (50 mM Tris-HCl, pH 7.5, 150 mM NaCl, 0.1% Tween-20) containing 5% skim milk powder for 1 hour and cut into sections to be probed with the indicated primary antibodies in the same buffer overnight at 4 °C. Membranes were washed six times with TBS-T then probed with horseradish peroxidase-conjugated secondary antibodies (Abcam) for 30 minutes at room temperature. After washing as above, membranes were incubated with enhanced chemiluminescence reagent (Millipore) and imaged with a MicroChemi chemiluminescence detector (DNR Bio-imaging Systems). The Gel Analyzer function on ImageJ was used to quantify protein signal.

**Purification of huntingtin-interacting proteins**

Cells were harvested by trypsinization, then washed with ice cold PBS. Proteins were crosslinked with 1% paraformaldehyde for 10 min followed by washing with PBS. Lysates were prepared in cytoskeleton (CSK) buffer (10 mM PIPES, pH 6.8, 100 mM NaCl, 300 mM sucrose, 3 mM $MgCl_2$, 1 mM EGTA, 0.5% (v/v) Triton X-100) containing 1X protein and phosSTOP inhibitor cocktails (Roche) for 5 min on ice and cleared by centrifugation. Supernatants were pre-cleared with rabbit IgG and Protein A agarose beads (MilliporeSigma) rotating for 30 min at 4°C. Pre-cleared products were collected and incubated in fresh Protein A agarose beads and anti-huntingtin EPR5526 (Abcam) for 2 hours rotating at 4°C. Immunoprecipitates were washed twice with CSK, twice with detergent-free CSK, and analyzed by mass spectrometry. For western blot analysis, RPE1 cells were treated with either HBSS or HBSS containing 400 μM $H_2O_2$ for 10 min and harvested as above.

**Mass spectrometry and protein identification**
MS/MS samples were analyzed using Sequest (XCorr Only) (Thermo Fisher Scientific, San Jose, CA, USA; version IseNode in Proteome Discoverer 2.2.0.388) and X! Tandem (The GPM, thegpm.org; version CYCLONE (2010.12.01.1)). Sequest (XCorr Only) was set up to search Uniprot-Mus-Oct252017.fasta (25074 entries) assuming the digestion enzyme trypsin. X! Tandem was set up to search a reverse concatenated Uniprot-Mus-Oct252017 database (50196 entries) also assuming trypsin. Scaffold (version Scaffold_4.8.4, Proteome Software Inc., Portland, OR) was used to validate MS/MS based peptide and protein identifications. Peptide identifications were accepted if they could be established at greater than 95.0% probability. Peptide Probabilities from X! Tandem and Sequest (XCorr Only) were assigned by the Scaffold Local FDR algorithm or the Peptide Prophet algorithm (99) with Scaffold delta-mass correction. Protein identifications were accepted if they could be established at greater than 20.0% probability and contained at least 1 identified peptide. Protein probabilities were assigned by the Protein Prophet algorithm(100). Proteins that contained similar peptides and could not be differentiated based on MS/MS analysis alone were grouped to satisfy the principles of parsimony.

**PAR overlay assay peptides**
PBM-1: LPRLQLELYKEIKKNGAPRSLR; PBM-2: KIIQLCDGIMASGRKAVTHAIPA; PBM-3: LLMCLIHFKSGMFRRITAAATRL; PBM-3 RR-AA: LLMCLIHFKSGMFAAITAAATRL; PBM-4: NKPLKALDTRFGRKLSIIRGIV.

**Protein purification**
Full-length PARP1 cDNA was subcloned into pFBOH-SBP-TEV-LIC and verified by sequencing. All proteins were produced in Sf9 culture as previously described (Harding et al, 2019). Briefly, Sf9 cells infected with P3 recombinant baculovirus were grown until viability dropped to 80–85%, normally after ~72 h post-infection, at which point cell pellets were harvested by centrifugation. HTT samples were purified by FLAG-affinity

chromatography and gel filtration as previously described (Harding et al, 2019). For PARP1, cell pellets were lysed in buffer containing 20 mM HEPES pH 7.4, 300 mM NaCl, 1 mM TCEP, 2.5% glycerol spiked with 1X protease inhibitor mix and clarified by centrifugation before affinity purification using TALON resin (Cytiva). PARP1-bound resin was washed with lysis buffer supplemented with 5 mM imidazole and then eluted with lysis buffer containing 250 mM imidazole. PARP1 samples were further purified by gel filtration using S200 16/60 column (Cytiva) equilibrated in 20 mM HEPES pH 7.4, 150 mM NaCl, 1 mM TCEP, 2.5% glycerol. The peak corresponding to monomeric PARP1 was pooled, concentrated ~10 mg/mL and aliquoted prior to flash-freezing in liquid nitrogen. All samples were assessed for purity by coomassie stained SDS-PAGE and were all shown to be >95% pure.
